# The regulatory architecture of the primed pluripotent cell state

**DOI:** 10.1101/2024.03.25.586622

**Authors:** Bo I. Li, Mariano J. Alvarez, Hui Zhao, Napon Chirathivat, Andrea Califano, Michael M. Shen

## Abstract

Although numerous studies have focused on defining transcriptional cell states in normal and disease contexts, the gene regulatory architecture that governs and defines specific mammalian cell states remains poorly understood. Here we present an integrative computational and experimental systems biology approach to elucidate the regulatory architecture of a conserved cell state of critical importance in development and stem cell biology, namely primed state pluripotency. We have used an unbiased approach to analyze protein activity profiles from mouse epiblast stem cells (EpiSCs), leading to identification and experimental confirmation of 132 transcription factors that are master regulators (MRs) of primed state pluripotency. These MRs include known as well as novel factors, many of which were further validated for their role in lineage-specific differentiation using CRISPR-mediated functional assays. To assemble a comprehensive regulatory network, we silenced each of the 132 MRs to assess their effects on the other MRs and their transcriptional targets, yielding a network of 1,273 MR→MR interactions. Network architecture analyses revealed four functionally distinct MR modules (communities), largely independent of lineage-specific differentiation, and identified key Speaker and Mediator MRs based on their hierarchical rank and centrality in mediating information flow in the pluripotent cell. Taken together, our findings elucidate the de-centralized logic of a “communal interaction” model in which the balanced activities of four MR communities maintain pluripotency, and define the primed pluripotent cell state in terms of its transcriptional regulatory network.

## Introduction

Pluripotency is defined by cellular states in which cells are capable of self-renewal while remaining poised to differentiate into all embryonic cell lineages. To date, a progression of three distinct *in vivo* pluripotency states—naïve, formative, and primed—has been identified in the mouse embryo^1–4^, with formative cells representing a transition between the naïve and primed states^5,6^. Due to their central significance for stem cell biology and regenerative medicine, the naïve and primed states have been captured as cell lines *in vitro* and intensively studied.

In culture, embryonic stem cells (ESCs) established from pre-implantation blastocysts reflect the naïve pluripotent state, whereas epiblast stem cells (EpiSCs) isolated from pre-gastrulation embryos^7,8^ correspond to the primed pluripotent state. Primed pluripotent cells co-express markers characteristic of the undifferentiated naïve state, as well as markers of more differentiated lineage-specific states; they differ from the naïve state in a range of molecular and functional properties^1,2,9^. Notably, conventional pluripotent stem cells established from human and many other mammalian species correspond to a primed pluripotency state, and thus most closely resemble mouse EpiSCs^1–3^. However, although the primed state is highly conserved, it is less understood than the naïve state, particularly in terms of the regulatory network that governs its pluripotency.

Although many genes have been implicated in pluripotency regulation, through functional studies on an individual gene basis, the regulatory logic and architecture that underlies the genetic control of pluripotency has remained elusive. Previous studies of pluripotency network architecture have largely focused on a limited number of well-known regulators that have been primarily identified by mutational analyses in cultured stem cells or *in vivo*^10–17^. These studies led to two general models of the regulatory logic underlying naïve state pluripotency. In the first model, a highly interconnected, hierarchical network of core transcription factors such as Oct4 and Nanog are responsible for activating pluripotency programs and repressing differentiation^10,11,13,17,18^. In the second model, pluripotency regulators comprise an array of specification genes that direct differentiation toward a specific lineage while repressing other lineages, creating an unstable undifferentiated equilibrium with the potential to commit to multiple lineages^19–23^. However, these models were based on analyses of a relatively small number of genes, and it is conceivable that key features that may have emerged from an unbiased approach were missed. More critically, elucidating large-scale network topology by studying a relatively small number of genes is suboptimal, as underscored by an ever-expanding repertoire of genes implicated in pluripotency maintenance, which suggests that pluripotency is controlled by a sizeable, highly complex gene regulatory network.

To address this challenge, we leveraged a comprehensive and unbiased systems approach for the large-scale systematic elucidation of the regulatory logic of EpiSC cells, a defined cell state of fundamental importance in normal development. The approach—which is based on the seamless integration of both computational and experimental methodologies—has led to the systematic identification and validation of primed pluripotency regulators, followed by elucidation of their regulatory relationships and overall network architecture. For this purpose, we have utilized ARACNe^24^ and VIPER^25^, two algorithms extensively validated in a wide range of physiologic^26-28^ and pathologic contexts, both tumor^29–32^ and non-tumor related^33–35^. ARACNe was used to assemble an initial regulatory network (interactome) from large collections of expression profiles, using an information theory approach that largely eliminates indirect interactions. VIPER interrogates the resulting interactome to identify candidate Master Regulator (MR) proteins representing mechanistic determinants of a specific transcriptional cell state via their transcriptional targets; this is accomplished by measuring the enrichment of their activated and repressed targets in genes differentially expressed in that state, akin to a highly multiplexed gene reporter assay. Notably, the accuracy and sensitivity of VIPER-inferred protein activity compares favorably with antibody-based measurements^35,36^, and allows unbiased identification of candidate MRs (*i.e.,* independent of prior literature knowledge).

In our study, we have generated an extensive collection of EpiSC gene expression profiles, as well as multiple signatures of lineage-specific differentiation to support the systematic generation of a mouse EpiSC-specific interactome and the identification of MR proteins responsible for the maintenance of primed state pluripotency. Using a set of 132 experimentally confirmed MRs, we elucidated their mutual regulatory relationships by systematically silencing each MR in turn to determine the resulting effects on the activity of all other MRs. This large-scale experimental analysis yielded a comprehensive MR→MR causal interaction network, which we have investigated using concepts from social network analysis to understand its regulatory architecture. Notably, topology analysis identified four distinct gene communities within the EpiSC network, suggesting a “communal interaction” model for how pluripotency is maintained. Our analysis of the regulatory architecture of primed state pluripotency thus provides novel insights into mechanisms of pluripotency and may guide future studies of other similarly complex regulatory networks.

## Results

### Overall experimental strategy

To dissect the logic of molecular interactions responsible for regulating primed state pluripotency, we pursued a six-step strategy that integrates computational inference with validation assays, followed by experimental refinement. First, as inputs for the ARACNe algorithm, we generated a large repertoire of gene expression profiles from EpiSCs undergoing a wide range of small molecule perturbations. This resulted in a comprehensive first-generation interactome comprising the computationally inferred transcriptional targets (*regulon*) of every transcription factor (Fig. 1a). Second, we leveraged this network to identify candidate MRs of primed state pluripotency, by using the VIPER algorithm to analyze gene expression time series representative of EpiSC differentiation along multiple lineages (Fig 1b). Third, we validated candidate MRs emerging from the analysis in terms of their ability to modulate the pluripotent transcriptional state following RNAi-mediated silencing, resulting in a list of 132 confirmed MRs that contained both known and novel regulators of pluripotency (Fig. 1c). Fourth, among these, we sought to validate 70 MRs not previously reported in the literature using functional assays of self-renewal and differentiation phenotypes following their CRISPR-mediated knock-out (Fig. 1d). Fifth, we used the RNA-seq profiles of EpiSC cells following RNAi-mediated silencing of each of the 132 confirmed MRs, as generated in step three, to assemble a comprehensive, experimentally-derived regulatory network governing primed state pluripotency (Fig. 1e). This fully causal MR→MR interaction network is based directly on experimental determination of the differential activity of each MR as induced by the silencing of every other MR. Finally, we investigated the architecture of the primed pluripotency network through modularity, hierarchy, and centrality analyses (Fig. 1f). Overall, this novel strategy allowed reconstructing the architecture of a complex and previously uncharacterized EpiSC regulatory network, using a fully unbiased approach that is largely based on the analysis of targeted experimental perturbations.

**Figure 1:**
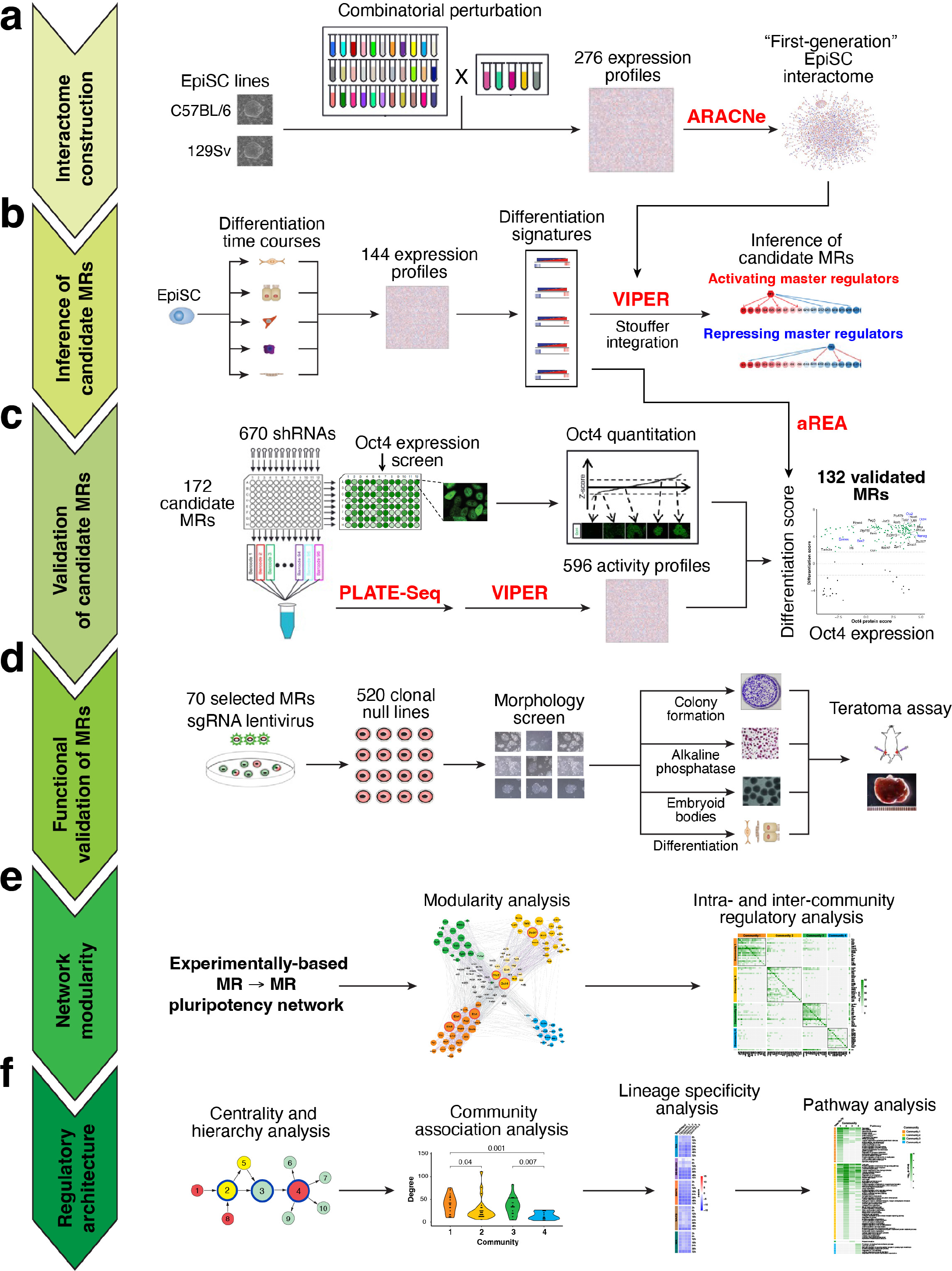
Flowchart of experimental design. See text for description.

### Computational inference of a primed state pluripotency network

The unbiased ARACNe-based^24^ reverse-engineering of a primed pluripotency cell state transcriptional interactome requires a large (*n*>100) compendium of gene expression profiles, with sufficient expression variability to reveal the range of mutual information between transcriptional regulators and their potential targets. To accomplish this goal, we developed a general strategy to create the necessary spectrum of EpiSC-derived expression profiles by combining two complementary sources of experimental perturbations. Specifically, we combined treatment with 33 small molecules (termed “perturbagens”) with 5 conditions known to induce differentiation. To be optimally informative and minimize underlying statistical co-dependencies, we selected perturbagens that affect an orthogonal range of signaling pathways to treat two distinct EpiSC cell lines. Critically, the order of treatment was defined based on a pilot experiment demonstrating greater variability of gene expression signatures when perturbagen treatment preceded induction of differentiation (Methods; Extended Data Fig. 1a; Supplementary Table 1).

This strategy generated 276 diverse gene expression profiles representing the response of the two EpiSC cell lines to combinatorial treatment with perturbagens and differentiation inducers (Methods; Extended Data Fig. 1a,b; Supplementary Tables 1, 2.1). These profiles were then analyzed by ARACNe to infer the transcriptional targets of 1,478 proteins annotated as transcription factors in the Gene Ontology^37^ database. The resulting EpiSC primed pluripotency interactome comprises 911,753 TF →target interactions between 1,393 TFs and 17,068 candidate transcriptional target genes. This first-generation interactome represents a foundational model for the subsequent identification of pluripotency maintenance master regulator (MR) proteins by VIPER analysis (Supplementary Table 3.1).

We then proceeded to identify candidate MRs for five distinct pathways of lineage-specific differentiation using the VIPER algorithm. VIPER requires signatures representing two distinct cellular states to identify the proteins (candidate MRs) representing mechanistic determinants of the transition between them. For this purpose, we generated gene expression profiles of EpiSCs following treatment with five distinct differentiation protocols at multiple time points (ranging from 3h to 72h). The resulting time series included a total of 144 gene expression profiles (Supplementary Table 2.2). These protocols were selected to enrich for differentiation along specific lineages, including neuroectoderm (RA and SB431542), trophectoderm (BMP4), mesoderm (WAFB), and endoderm (WA) (Methods; Extended Data Fig. 1c-f). To confirm that these differentiation treatments induced distinct transcriptional states, we analyzed the resulting time series by Principal Components Analysis (PCA) and by expression of lineage-specific marker proteins as assessed by immunofluorescence (Extended Data Fig. 1c-f). For each time point in this analysis, we generated a differential gene expression signature for VIPER analysis by comparing perturbagen-treated to baseline untreated EpiSC cells. The resulting differentiation signatures were then analyzed by VIPER, using the EpiSC interactome, to identify candidate MRs for the associated cell state transitions.

We reasoned that the most universal pluripotency MRs would be those conserved across most if not all of the five differentiation pathways examined. We thus used Fisher’s method to integrate the VIPER statistics (p-values) representing the differential activity of 1,393 TFs, as determined by the analysis of each differentiation signature, across all time points and differentiation trajectories. The resulting ranked list identified 300 candidate MRs whose regulons were significantly enriched in differentially expressed genes (p < 10^-4^), with negative and positive enrichment representing MRs capable of inducing or repressing pluripotency, respectively (Fig. 2a; Supplementary Table 4.1). From this initial list, we selected 172 candidate MRs for further analysis, including most of those that were top ranked, as well as additional candidate MRs of interest (see Methods for selection criteria). Many of the candidate MRs nominated by this systematic approach had been previously implicated in pluripotency regulation (Supplementary Table 4.4), confirming that our unbiased analysis was highly effective and suggesting that the remaining novel candidates might include many *bona fide* pluripotency MRs.

**Figure 2.**
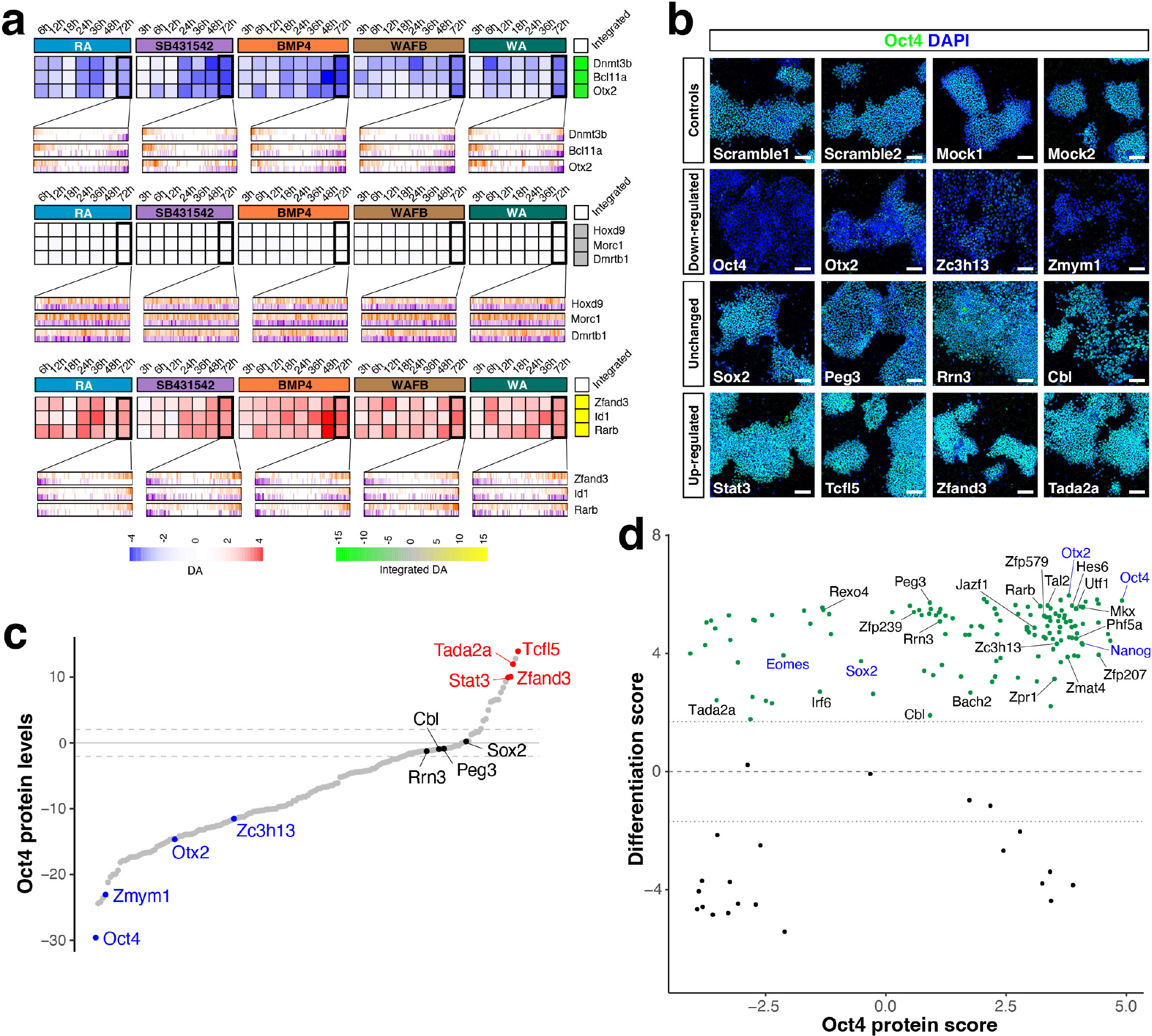
Identification and validation of candidate master regulators of primed state pluripotency. (a) Examples of transcriptional regulatory proteins being inactivated, unaffected, or activated during the differentiation treatments. Each heatmap cell represents the VIPER-inferred differential protein activity expressed as normalized Enrichment Score (NES). Enrichment plots show the distribution of activated (orange) and repressed (purple) transcriptional targets on the 72h gene expression signatures. The right-most column represents the integrated differential activity of each protein across all time points and treatments. RA: Retinoic acid; WAFB: Wnt3a, Activin A, FGF2, and BMP4; WA: Wnt3a and Activin A. (b) Oct4 staining of EpiSCs after shRNA-mediated silencing of selected candidate MRs and vehicle or mock-treated controls. Scale bars: 100 microns. (c) Waterfall plot showing differential Oct4 protein level after silencing of each candidate MR; dashed lines indicate FDR < 0.05 (2-tailed). Raw data are provided in Supplementary Table 5.2. (d) Scatterplot of the effect of silencing each MR on Oct4 protein level and the differentiation score of EpiSCs, which is determined by the recapitulation of the protein activity signature of EpiSC differentiation by the silencing experiment (see Methods). We observed a significant correlation between these scores (Spearman’s Rho 0.32, p < 10^-4^); dotted lines indicate FDR = 0.05 (2-tailed). Candidate MRs showing significant validation (differentiation) score are highlighted in green. Selected MRs labeled in black indicate MRs functionally validated in CRISPR/Cas9 assays, and in blue indicate known key pluripotency regulators. Raw data are provided in Supplementary Table 6.2.

### Experimental confirmation of candidate MRs

To systematically confirm the functional role of the 172 candidate MRs in pluripotency control, we employed two independent approaches. First, we assessed whether lentiviral-mediated shRNA silencing of candidate MRs would modulate the expression of endogenous Oct4 (Pou5f1) at the protein level (Fig. 2b,c). To minimize contributions from off-target effects, we used multiple independent hairpins for each candidate MR, with 92% of the candidate MRs targeted by at least 3 hairpins (Extended Data Fig. 1g; Supplementary Table 5.1). We quantified Oct4 protein levels by immunofluorescence staining, and calculated z-scores representing its differential protein expression relative to non-targeting controls (Fig. 2c). Using this assay, we found that 117 (68%) and 15 (8.7%) of the candidate MRs down-regulated and up-regulated Oct4 protein expression when silenced, respectively (FDR < 0.05) (Fig. 2c; Supplementary Table 5.2). Notably, the analysis identified Oct4 itself as well as other well-known MRs such as Otx2 and Stat3^38,39^ as pluripotency regulators (Fig. 2b,c).

Secondly, to confirm the roles of candidate MRs that modulate pluripotency independently of their influence on Oct4 protein expression, we developed an assay to determine whether their silencing might recapitulate the EpiSC differentiation signature. This assay utilized PLATE-seq, a multiplexed barcoded RNA-sequencing protocol^40^ that generates low-depth (1∼2 million reads/sample) gene expression profiles. We used PLATE-seq to generate gene expression profiles produced by 596 independent shRNA-mediated silencing assays that targeted 154 of the 172 candidate MRs in EpiSC cells; the other 18 candidate MRs were either targeted unsuccessfully or lacked a regulon for analysis (Methods; Supplementary Table 5.1). Next, we performed VIPER-based analysis of the PLATE-seq profiles, using the EpiSC interactome, to assess the differential activity of all candidate pluripotency MRs, as induced by each silencing assay vs. the pool of non-targeting controls. The resulting differential MR activity profiles were compared to those obtained along the five lineage-specific differentiation time courses by enrichment analysis using the aREA algorithm^25^. The resulting Normalized Enrichment Scores (NES) from all hairpins targeting the same candidate MR were integrated across all five differentiation signatures, using a weighted implementation of Stouffer’s method. Weights were proportional to the effect of each hairpin on the activity of its target protein, such that hairpins inducing less-efficient silencing had lower weight (Supplementary Table 6.1).

Interestingly, we have previously shown that up to ∼20% of candidate VIPER-inferred MRs may be correctly inferred but with an opposite directionality (inverted NES sign), due to the presence of autoregulatory loops that invert the relationship between mRNA expression and the activity of its associated protein at steady state (equilibrium)^25^. As a result, these analyses were instrumental in confirming the role of candidate MRs as either activators or repressors. Furthermore, we also identified and corrected the activity of 27 of the tested MRs (18%) that had been inferred with inverted activity (Methods; Supplementary Table 6.2).

To determine the number of candidate MRs whose induced differentiation signature was consistent with or divergent from their ability to modulate Oct4 protein expression, we computed a Marker Score (MS) and a Differentiation Score (DS; see Methods for details) (Fig. 2d; Supplementary Table 6.1). As expected, the two scores were highly correlated (*p* = 4.8 x 10^-5^), with 85 of 154 candidate MRs tested by both assays (55%, p < 0.05, 1-tailed FET) showing concordant scores (FDR < 0.05, by 2-tail analysis). Notably, an additional 47 candidate MRs (31%) presented with a significant DS but not MS score, suggesting that these genes may regulate primed state pluripotency in a Oct4-independent manner. In total, we confirmed 132 MRs (86%) by the more conservative test (DS analysis, green dots in Fig. 2d), which were selected as the basis for further investigation.

### Differentiation analysis of novel primed state pluripotency MRs

Since the ability of an MR to induce the experimentally determined differentiation signature upon silencing suggests a potential mechanistic role in pluripotency control, we tested whether silencing of each confirmed MR was sufficient to induce differentiation in culture, using several cell-based assays. For this purpose, we focused on 70 of the 132 confirmed MRs that were identified as novel (*i.e.,* not previously reported in the pluripotency literature at the time the list was compiled), as well as on 4 additional genes (Utf1, Hes6, Rarb, and Ilf3) selected from the MRs previously implicated in pluripotency regulation as positive controls (Supplementary Table 4.2). For functional validation, we performed CRISPR-mediated gene knock-out assays (CRISPR-KO) using a lentiviral system^41^ to generate mutant alleles for each MR, thus yielding 1,150 clonal EpiSC lines and 128 scramble or mock control lines (Supplementary Table 7.1). Of the 1,150 clonal lines, 311 contained monoallelic frameshift mutations, while 319 contained biallelic in-frame mutations that may not be loss-of-function. The remaining 520 lines contained biallelic frameshift mutations and were thus identified as candidate null mutants for 62 of the targeted novel MRs and the 4 selected known MRs. We failed to derive candidate null lines for the other 8 novel MRs, 3 of which (Eno1b, Zfp598, and Zfp184) might be due to the low number of clonal sgRNA-targeted lines obtained (Supplementary Table 7.1). However, of the other 5 remaining MRs (Tada2a, Phf5a, Zpr1, Zfp207 and Rrn3), we derived ≥ 20 clonal targeted lines for each, none of which represented a biallelic frameshift mutant, suggesting that null mutants for these genes may be impaired in self-renewal and/or clonogenicity (*p* < 0.01 by Chi-square test; Extended Data Fig. 2a).

We screened the candidate null mutant and control targeted lines for their morphological phenotypes in feeder-free culture conditions, in which wild-type EpiSC cells display round and densely packed colonies with large nuclei, prominent nucleoli, and scant cytoplasm. We found that candidate null mutants for 15 MRs displayed a higher frequency of abnormal morphologies relative to controls (Fig. 3a, Extended Data Figs. 3 and 4a). In addition, shRNA-mediated knock- down of the 5 MRs that failed to generate biallelic frameshift mutants also resulted in abnormal morphologies (Extended Data Figs. 3 and 4a). Overall, 3 of the 4 (75%) MRs previously implicated in pluripotency regulation as well as 17 of the 70 (24%) novel MRs displayed morphological phenotypes in this analysis, demonstrating its robustness in identifying pluripotency regulators validated by multiple assays. For further analyses of these 20 MRs, we selected a total of 40 knock- out (KO) lines for the 15 MRs with biallelic frameshift mutations and 11 knock-down (KD) lines for the remaining 5 MRs, as well as 9 negative controls (Extended Data Fig. 2b; Supplementary Table 7). To assess whether these KO/KD lines were truly null, we performed qRT-PCR analyses of targeted MR expression, which showed significantly lower expression in 46 of the 51 KO/KD lines (Extended Data Fig. 5), consistent with increased RNA degradation induced by frameshift mutations or RNA interference.

**Figure 3.**
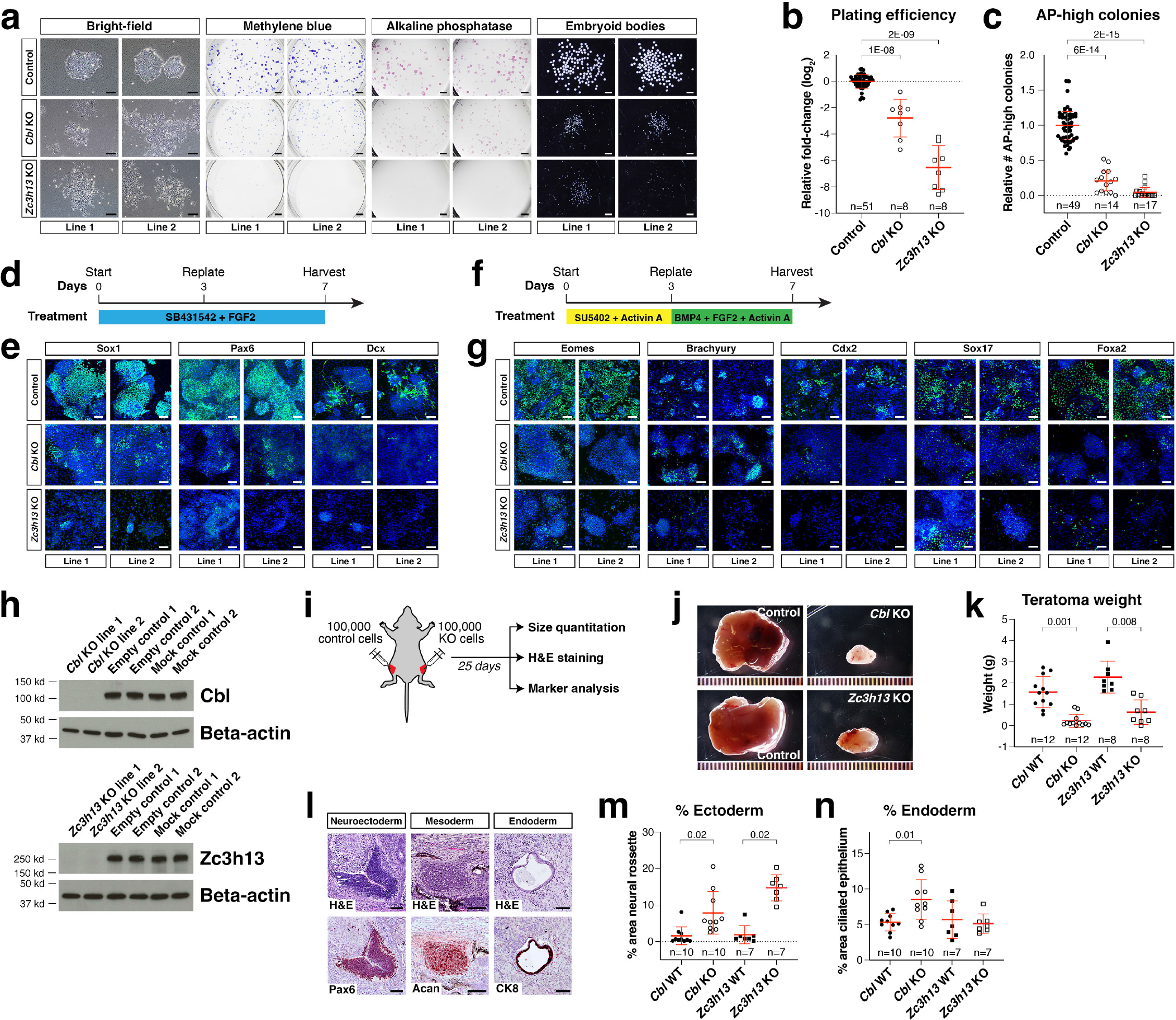
Functional validation of two master regulators. (a) Morphology of KO or control EpiSC lines; representative images for independent lines are shown. (b) Fold change in cell number after five days of colony formation. (c) Number of alkaline phosphatase positive colonies. (d) Time line of neuroectoderm differentiation. (e) Immunofluorescence staining of biological replicates (clonal lines) for lineage-specific markers. (f) Time line of mesendoderm differentiation. (g) Staining for lineage-specific markers. (h) Western-blot for Cbl and Zc3h13. (i) Schematic of the teratoma assay. (j) Representative images of mutant and control teratomas dissected from contralateral legs of the same animal. (k) Teratoma weight. (l) Hematoxylin and eosin (H&E) and immunohistochemical staining of serial sections. (m, n) Fraction of area showing ectoderm (neural rosette) (m) and endoderm (ciliated epithelium) features (n). Line and whiskers show the mean ± s.d.; statistical significance is shown as FDR, 2-tailed U-test. Scale bars: 100 microns.

We analyzed the self-renewal and differentiation potential of the 20 MRs using *in vitro* functional assays. We found that KO/KD lines for 18 of the 20 MRs presented lower colony formation efficiency, with 13 of these displaying significantly fewer alkaline phosphatase (AP)- high colonies (Fig. 3a-c, Extended Data Figs. 3 and 4a-c; 2-tailed U-test; FDR < 0.05). We used embryoid body (EB) formation to assess pluripotency exit, and found significantly reduced EB formation efficiency in KO/KD lines for 18 MRs (Fig. 3a, Extended Data Figs. 3 and 4a). Finally, we performed directed differentiation to neuroectoderm or mesendoderm followed by immunostaining for lineage-specific markers, and observed significant effects on neuroectoderm differentiation in 16 KO/KD lines, and on mesendoderm differentiation in 8 KO/KD lines (Fig. 3d-g, Extended Data Figs. 3 and 4a,d,e). Interestingly, none of the 20 functionally validated MRs displayed similar effects across all assays, indicating the functional diversity of individual MRs in regulating distinct aspects of pluripotency and lineage differentiation, as well as the robustness of our MR discovery pipeline. Finally, during the time span of our experiments, 6 of the 20 validated MRs (Irf6, Jazf1, Peg3, Phf5a, Zc3h13, and Zfp207) were identified as pluripotency relevant, albeit in non-EpiSC contexts^42–47^, providing independent confirmation of our systematic MR analyses.

We focused in greater detail on 2 MRs that consistently generated highly significant results in nearly all assays, Cbl and Zc3h13. Cbl is a member of a family of three closely related RING finger E3 ubiquitin ligases that are responsible for proteasome-mediated substrate degradation^48^; although *Cbl* null mutant mice are homozygous viable with T cell defects, *Cbl; Cbl-b* double homozygotes display an uncharacterized lethality prior to mid-gestation^49^. Zc3h13 is a component of the WMM (Wtap-Mettl3-Mettl14) complex that mediates N6-methyladenosine (m6A) methylation of RNA, and its knock-down decreases self-renewal of mouse embryonic stem cells^43^; in addition, we have found that *Zc3h13* null mutants display a peri-implantation lethal phenotype (N. Chirathivat *et al.,* in preparation). We confirmed that the KO EpiSC lines for *Cbl* and *Zc3h13* were true null alleles by Western blotting (Fig. 3h), and that *Cbl* and *Zc3h13* mutant EpiSCs displayed altered differentiation of neuroectoderm and mesendoderm at the mRNA level by qRT-PCR (Extended Data Fig. 6). To determine whether loss-of-function for *Cbl* and *Zc3h13* affected differentiation potential *in vivo*, we performed grafts of KO lines to generate teratomas and found that both *Cbl* and *Zc3h13* KO teratomas were smaller in size and weight (Fig. 3i-k). By H&E staining and marker immunostaining of tissue sections, we found that *Cbl* and *Zc3h13* KO teratomas displayed increased neural rosette formation and neural marker expression, whereas *Cbl* KO but not *Zc3h13* KO resulted in increased ciliated epithelium, consistent with increased epithelial marker expression (Fig. 3l-n, Extended Data Fig. 7). Thus, both Cbl and Zc3h13 are required for pluripotency maintenance in teratomas.

### An experimentally-based primed state pluripotency network

Having computationally identified a large number of previously known as well as unknown MRs for primed state pluripotency, and functionally validated many of the previously unknown ones, we next sought to elucidate the architecture of the regulatory network that governs the primed state pluripotency state. To accomplish this goal, we derived an experimentally-refined, second-generation primed pluripotency transcriptional interactome, by leveraging the PLATE-seq gene expression profiles representing the response of EpiSC cells to the silencing of each of the 132 confirmed MRs (green dots in Fig. 2d). Specifically, these data were used to refine the ARACNe regulon of each confirmed MR by integrating the statistical significance of *(a)* the candidate target’s differential expression following MR silencing and *(b)* their mutual information. By representing the causal response of the cell to silencing of each MR, this strategy also helped infer the directionality of each interaction. This analysis produced a comprehensive set of experimentally refined regulons, resulting in an “experimentally curated,” causal interactome (ECC-interactome) (Supplementary Table 3.2).

Using the ECC-interactome, we evaluated whether silencing a specific MR would affect the VIPER-inferred activity of another MR, as assessed using the experimentally refined regulons (Supplementary Table 3.2). This analysis yielded a pluripotency focused, experimentally-based EpiSC regulatory MR network (Fig. 4a), containing 1,273 MR→MR interactions (edges) between 120 of the 132 confirmed MRs (FDR < 10^-10^) (Fig. 4a and Supplementary Table 8).

**Figure 4.**
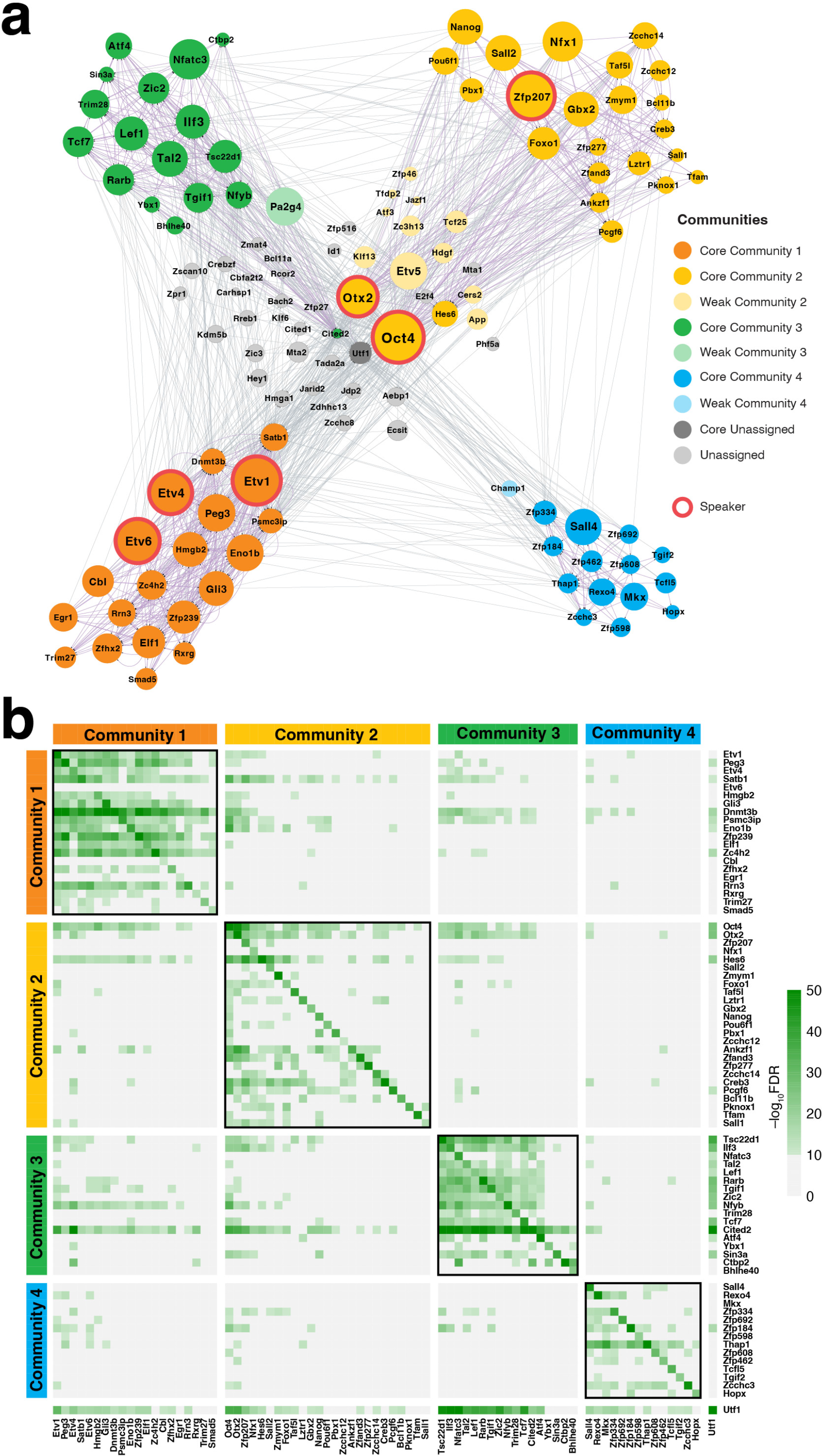
Regulatory network model for primed state pluripotency. (a) Regulatory network of primed state pluripotency with 120 nodes and 1,273 unidirectional edges. Only significant effects (FDR < 10^-10^, 2-tailed aREA test) are included. The size of each node is proportional to its Regularized Out-Degree (ROD) score; colors indicate community assignment (strong, FDR < 0.01; weak, 0.01< FDR < 0.1; unassigned FDR> 0.1). Six outlier MRs showing the highest ROD scores (Speakers) are highlighted. (b) Heatmap showing the effect of silencing each MR on the activity of all other MRs. 77 MRs (76 strong community members and Utf1) are grouped by their community membership; color scale indicates statistical significance (-log_10_(FDR), 2-tailed aREA test) for each MR protein differential activity.

To investigate the architecture of this EpiSC pluripotency MR network, we assessed its modularity, using Louvain’s community detection algorithm^50^. The analysis showed that most of the confirmed MRs were organized into four distinct communities (Extended Data Fig. 8a). We then assessed the association between each MR and each community, based on the statistical significance of the enrichment of the MRs representing its first neighbors (*i.e.,* the MRs directly connected to it) in each community. The analysis identified 76 MRs with strong community association (FDR < 0.01, 1-tail Fisher’s Exact Test, FET), 13 MRs with weak association (0.01 ≤ FDR < 0.1), and 31 MRs lacking association (FDR ≥ 0.1) with any community (Fig. 4b; Supplementary Table 8.4; Extended Data Fig. 8a).

### The regulatory logic of primed state pluripotency

We next assessed whether individual MRs may play distinct roles in the EpiSC pluripotency MR network, based on their interactions with other MRs. For this purpose, we assessed the In-Degree and Out-Degree metrics for each MR, corresponding to its number of incoming (→MR) and outgoing (MR→) regulatory interactions; we also computed a “Regularized Out-Degree” (ROD) metric, which corrects for the different number of interactions of each MR (see Methods). The statistical significance of their In-Degree vs. Out-Degree helped stratify MRs into three classes, corresponding to “Speakers” (significantly higher Out-Degree), “Listeners” (significantly higher In-Degree), and “Communicators” (no significant difference) (Fig. 4a; Fig. 5a-c), thus illuminating their role in mediating the flow of information in pluripotent cells. We could also identify “Mediator” MRs based on their “Betweenness” centrality, a metric aimed at assessing the number of shortest paths between any two MRs going through an MR of interest, thus capturing its influence over the flow of information in the network (Fig. 5a,c; Extended Data Fig. 9a,b). Importantly, for the 77 Core MRs (76 with strong community association plus Utf1), there was no correlation of their average protein activity, community score, ROD, or Betweenness with the silencing efficiency of shRNA hairpins used in our PLATE-seq analysis (Rho < 0.22, p > 0.01; Extended Data Fig. 9c-f), suggesting the absence of assay-derived bias. However, we did observe ROD differences between communities 1 and 4, as well as differences in interaction degree between communities (Fig. 5d-f), suggesting differences in the connectivity of these communities.

**Figure 5.**
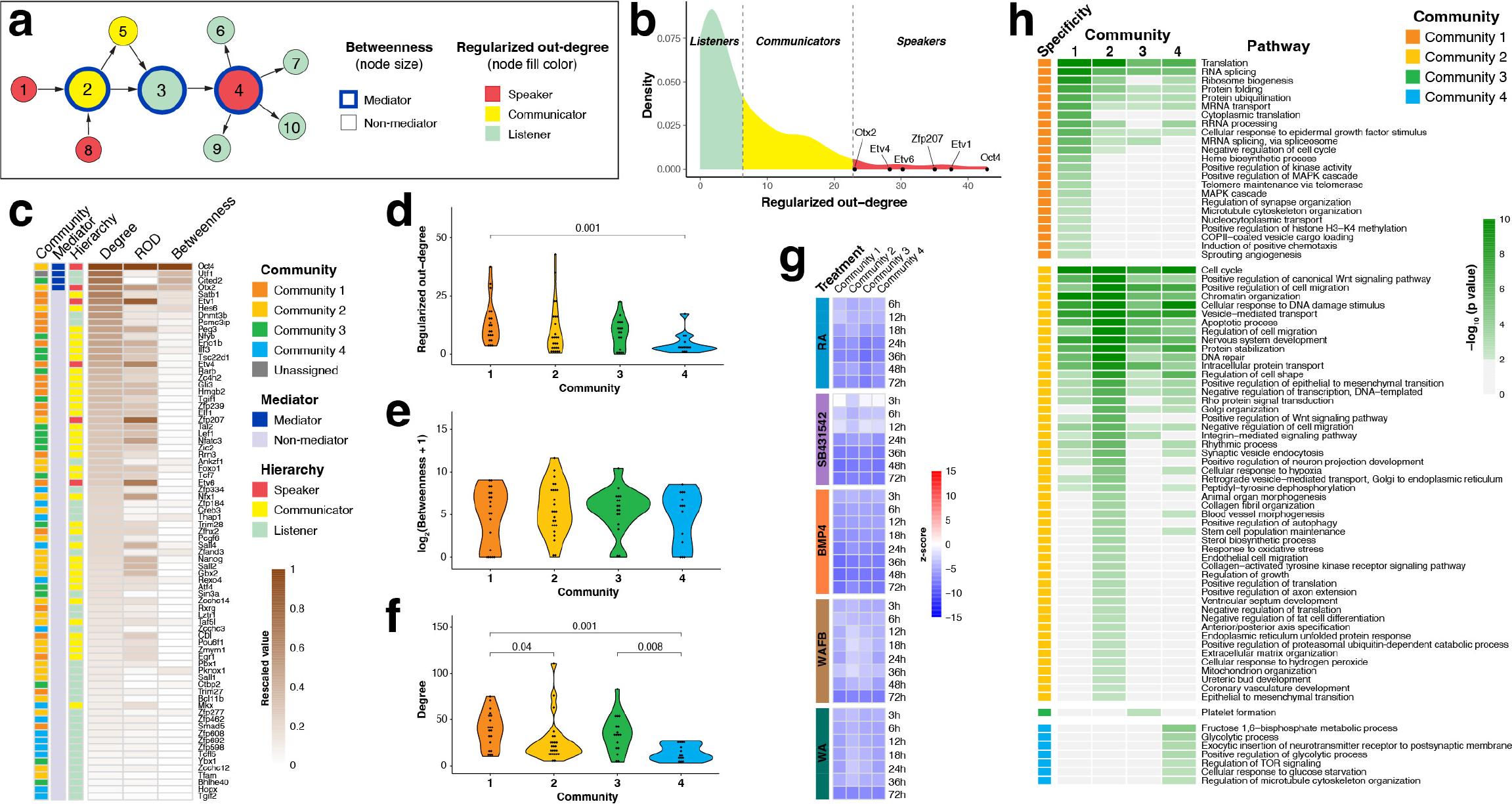
Community structure of the primed state pluripotency network. (a) Schematic representation of parameters for defining node hierarchy based on their regularized out-degree (ROD) and betweenness score. (b) Distribution density for ROD across all MRs. The vertical dotted lines indicate the point of equal probability for a mixture of gaussian models fitted to the data, defining the boundaries for Listeners, Communicators and Speakers. (c) Heatmap showing network parameters for the 77 MRs, showing scaled values for Degree, ROD and betweenness scores. (d-f) Violin plots showing distribution of regularized out-degree (d), betweenness (e), and degree (f) (FDR < 0.05, 2-tailed U-test). (g) Heatmap of the integrated meta-protein activity change for each community across the five differentiation treatments. Each cell represents the weighted average of the relative change in activity vs. untreated EpiSC controls across all MRs of each community, with the weight calculated based on the MR community score (Extended Data Fig. 8). (h) Heatmap showing GOBP pathways preferentially enriched (-log_10_(p-value)) on each of the four communities.

These analyses identified 6 Speaker, 75 Listener, and 39 Communicator MRs, as well as 4 Mediator MRs (Oct4, Cited2, Utf1, and Otx2) (Figs. 4a, 5a,c; Extended Data Fig. 9b). The Speakers include three ETS transcription factors (Etv1, Etv6, and Etv4), likely due to their roles in mediating FGF signaling, which is involved in maintaining primed state pluripotency^7^. Interestingly, Zfp207, which has only recently been characterized in regulation of pluripotency^44,51^, was identified as a Speaker, while Utf1 was the only Speaker or Mediator not strongly associated with any of the four communities. Notably, two well-characterized pluripotency regulators, Oct4 and Otx2, were identified as both Speakers and Mediators (Fig. 4a; Extended Data Fig. 9b; Supplementary Table 8.2), reinforcing their significance as key pluripotency regulators.

Next, we asked whether the arrangement of MRs into communities might reflect differences in their functions in pluripotency maintenance or lineage specification. Therefore, we examined the differential activity of each MR in the network along the five lineage differentiation time courses. Interestingly, nearly all MRs displayed similar patterns, with ∼10% and ∼90% of the MRs showing time-dependent increased and decreased activity along the five differentiation treatments, respectively, suggesting a highly coordinated program (Extended Data Fig. 8b). To determine whether individual communities may be associated with specific lineage differentiation paths, we then integrated the activity of the MRs in each community weighted by their community scores (Extended Data Fig. 8a). Interestingly, the average activity of all four communities was significantly reduced across all five lineage differentiation signatures (Fig. 5g), indicating that the four communities are not associated with any specific lineage differentiation trajectory.

Finally, to determine whether individual communities oversee distinct biological functions, we performed pathway enrichment analyses on the gene expression signature obtained by silencing of each individual MR. We then integrated the enrichment scores for the MRs assigned to each community while weighting their contribution by the community association scores (Fig. 5h and Extended Data Fig. 8a). This analysis identified 250 Gene Ontology-Biological Process (GOBP) terms enriched in at least one community, with 57 enriched in all four (FDR < 0.01, 1-tailed aREA test; Supplementary Table 9.1). Common biological terms included “developmental processes,” “nucleotide processing,” “protein modification,” and “cellular transportation events,” all of which are involved in pluripotency regulation. In contrast, 81 GOBP terms were preferentially enriched only in a single community (Fig. 5h; Supplementary Table 9.2). Community 1, for instance, is enriched in “nucleotide processing,” “protein modification,” and “other intracellular activities,” Community 2 in a range of developmental processes and cellular signaling pathways, and Community 4 in metabolic pathways. Interestingly, Community 3 was only associated with a single category, “platelet formation,” thus defying a more general characterization. Taken together, pathway enrichment analysis revealed both cooperative and unique roles of MR communities in regulating primed state pluripotency.

## Discussion

Pluripotent cell states and dynamic transitions between these and more differentiated states have long been utilized as model systems for reconstruction of gene regulatory networks. For example, development of the CellNet platform revealed a regulatory network for somatic cell reprogramming and identified factors modulating the efficacy of these cell fate conversions^52–55^. However, earlier studies of pluripotency networks have focused only on a relatively small set of regulators, many of which had been previously identified in the literature. Our current network for primed state pluripotency is notable for its size, unbiased generation, and derivation by computation followed by direct experimentation. Furthermore, the size of our network has allowed analysis of its overall topology and elucidation of its regulatory logic.

Our analysis includes several advances that taken together produce an entirely novel and highly generalizable methodology. Specifically: (*a*) Nodes (*i.e.,* MRs) in the regulatory network were identified by an unbiased methodology that is not based on prior knowledge of gene function; (*b*) Node identification was not dependent upon differential expression of the corresponding gene, but rather on the role of the encoded protein in regulating the overall transcriptional state of the pluripotent cell (*differential activity*), as determined by the differential expression of its transcriptional targets. This is critical since transcription factor activity is often regulated post-translationally rather than transcriptionally; (*c*) Network interactions (i.e., edges) and causality were dissected based on the effect of regulatory proteins on the inferred activity of other regulatory proteins, following experimental silencing of 132 confirmed MRs identified by the analysis; (*d*) Assessment of pluripotency integrated the effect of individual regulators on Oct4 protein expression as well as on induction of lineage-specific differentiation signatures, providing an approach to identify both Oct-4 dependent and Oct4-independent pluripotency regulators; (*e*) Analyses of network architecture utilized concepts of centrality and betweenness, which are critical properties of social network interactions^56,57^. This approach is instrumental in illuminating the control of information flow in primed pluripotent cells, based on the well-studied concept in graph theory that some nodes may play a critical role in mediating the information flow between communities (Mediators) or in controlling entire programs (Speakers)^56^. In combination, these features result in an unbiased primed state pluripotency network architecture that is substantially larger than described in previous studies.

We have shown that 76 of the 120 regulatory proteins that control pluripotency in our network are organized within four regulatory modules, termed “communities,” where the highly interconnected architecture of each community is countered by a much sparser connectivity between communities. This network architecture contrasts significantly with the previously hypothesized “hierarchy” and “balance” models. Instead, our findings suggest that, in mouse primed state pluripotent cells, the “hierarchy” and “balance” models coexist in a hybrid form that we term “communal interaction”. Specifically, we propose that primed state pluripotency is maintained by the balanced activities of four regulatory communities, with a small number of community members providing hubs for information flow. Each community represents a distinct functional unit yet is not closely associated with lineage-specific differentiation, as originally proposed in the balance model.

In principle, identification of 8 MR proteins as either Speakers or Mediators might lend support to the existence of core factors in the network. Notably, Mediators and Speakers include Oct4 and Otx2, consistent with their essential roles in primed state pluripotency^38,58,59^. In addition, the identification of ETS transcription factors Etv1, Etv4, and Etv6 as Speakers is consistent with the central role of FGF signaling in maintaining primed state pluripotency^7^. However, previous studies have not suggested a critical role for Zfp207, Utf1, and Cited2 in controlling primed state pluripotency, suggesting that these proteins may deserve further investigation. Importantly, EpiSCs differ from mESCs in their lack of requirement for key regulators of the naïve pluripotent state such as Sox2 and Nanog^60,61^, and notably these genes were not detected as having central roles. Overall, however, the strong association of 7 of the 8 Speaker/Mediator MRs with one of the 4 communities argues against a dominant role of these MRs. Rather, the nearly equivalent levels of hierarchical organization and centrality among the four communities (Fig. 5d-f) supports a more decentralized architecture of the primed state pluripotency network.

It is important to note some limitations of the approach that may affect the completeness and accuracy of our regulatory network for primed state pluripotency. First, the analysis is limited to 1,393 genes annotated as transcription factors. Second, MRs are often synergistic, even if they fail to produce an observable phenotype on an individual basis^29,30^; consequently some of the MRs identified by this study may lack a detectable phenotype *in vitro* and/or *in vivo*, suggesting that the number of MRs contributing to primed pluripotency maintenance may be larger than that validated by our functional assays. Finally, this study was performed using bulk, rather than single-cell profiles, and consequently may not account for potential heterogeneity within EpiSC cultures.

More broadly, our work provides a general road-map for elucidating the regulatory logic of a complex cell state. In particular, when combined with gene expression and protein activity analysis, our methodology for low-cost, high-throughput RNA-sequencing facilitates practical direct experimentation to infer or refine the architecture of any regulatory network. Furthermore, our study provides a rich and valuable resource to investigate molecular mechanisms that govern stemness. Future studies will promote development of innovative strategies to control the dynamics of pluripotency and may therefore hold considerable promise for translational applications in regenerative medicine.

## Acknowledgements

We thank Jianhua Chu for generating and providing the EpiSC-G line, Ron McKay for generously providing the EpiSC-5 line, Emlyn Parfitt for advice on perturbagen concentrations, Forest Ray, Erin Bush, and Peter Sims for advice on PLATE-Seq, and Cory Abate-Shen and Jianlong Wang for comments on the manuscript. Imaging using the Citation 5 Indigo instrument was performed in the Confocal and Specialized Microscopy Shared Resource of the Herbert Irving Comprehensive Cancer Center at Columbia University Irving Medical Center, supported by NIH/NCI Cancer Center Support Grant P30 CA013696. This work was supported by NIH R01 HD085904 to M.M.S. and by an NCI Outstanding Investigator Award (R35 CA197745) and NIH Shared Instrumentation Grants S10 OD012351, S10 OD021764, and S10 OD032433 to AC.

## Author contributions

Conceptualization: B.I.L., M.J.A., H.Z., A.C., and M.M.S.; Methodology: B.I.L. and M.J.A.; Software: M.A.; Formal analysis: B.I.L, M.J.A., and A.C.; Investigation: B.I.L., H.Z., and N.C.; Data curation: M.J.A.; Writing (original draft): B.I.L. and M.J.A.; Writing (review and editing): B.I.L., M.J.A., A.C., and M.M.S.; Visualization: B.I.L., M.J.A., A.C., and M.M.S.; Supervision: A.C. and M.M.S.; Funding acquisition: A.C. and M.M.S.

## Materials and Methods

### EpiSC derivation and maintenance

All experiments utilized the EpiSC-5 line^8^, which was derived from 129SvEv mice, or the EpiSC-G line. We derived the EpiSC-G line from C57BL/6 mice as described^7,8^, with the following minor modifications. In brief, C57BL/6 pregnant female mice were euthanized at 5.5 dpc, and pre-gastrulation epiblast was dissected intact and separated from extraembryonic ectoderm using glass needles in calcium- and magnesium-free PBS. Dissected tissue was treated with Cell Dissociation Buffer (Gibco 13151-014) at 4°C for 15 min. Epiblast cells were cultured on inactivated mouse embryonic fibroblasts (MEFs) in EpiSC basal medium containing DMEM/F12, 1x GlutaMAX supplement (Gibco 10565-018), 20% knockout serum replacement (Gibco 10828-010), 1x MEM non-essential amino acids (Gibco 11140-050), 0.1 mM 2- Mercaptoethanol (Gibco 21985-023), 10 ng/ml Activin (Gemini 300-356P), and 12.5 ng/ml FGF2 (Peprotech 100-18B). Every 3-4 days, cells were dissociated with 2.5 U/ml Dispase (Stemcell Technology 07913), and re-plated as small clumps at ∼1:6 ratio onto a new MEF layer. After 8∼10 passages, cells were dissociated, aliquoted, and frozen in cryovials in 50% basal medium/40% knockout serum replacement/10% DMSO. The established cell line was verified for normal karyotype and tested negative for mycoplasma.

EpiSC lines were cultured under feeder-free conditions in Matrigel-coated plates with filtered MEF-conditioned media (EpiSC basal media harvested after culturing with inactivated MEFs for 24 h) supplemented with 12.5 ng/ml FGF2 and 10 ng/ml Activin (CMAF). In experiments requiring cell quantitation, cells were dissociated with 0.05% Trypsin or TrypLE and counted with an automated cell counter (Bio-Rad TC20). Cells were then re-plated onto Matrigel-coated plates with CMAF supplemented with 10 uM Rock inhibitor Y-27632.

### EpiSC differentiation for interactome generation and analysis

To generate gene expression profiles for interactome assembly, we used the EpiSC-5^8^ and EpiSC-G lines. Cells were plated at a density of 50,000 cells/well in 6-well plates coated with Matrigel and 2 ml CMAF + 10 µM Y27632. At 24h after plating, cells were maintained in CMAF or switched to EpiSC basal media (Supplementary Table 2.1), followed by addition of each of 33 perturbagens (Supplementary Table 1). Each one of five differentiation-inducing treatments (Supplementary Table 1) was then added at 12h after perturbagen addition; finally, cells were harvested at 60h after initial plating. For embryoid body formation, perturbagen-treated EpiSCs were dissociated and plated onto AggreWell^TM^ 400 plates (StemCell Technology 34411), with 2 ml of EpiSC basal media at 1.2 million cells/well. Plates were spun for 5 min at 1500 rpm to settle the cells. At 24h after plating, the nascent embryoid bodies were transferred to 6-well, low-attachment plate cultures (Corning 3471) for an additional 12h before harvest. Gene expression profiles were generated as described in the transcriptomic profiling section.

Differentiation time course data were generated using the EpiSC-5 line. For retinoic acid treatment, cells were cultured with 5 µM retinoic acid in MEF conditioned media without supplementation of FGF2 or Activin A (CM) for 72h. For BMP4 treatment, cells were cultured for 72h in CM + 100 ng/ml BMP4. For SB431542 treatment, cells were cultured for 72h with 10 nM SB431542 in culture media containing DMEM/F12 with 1x GlutaMAX/1x insulin-transferrin-selenium/1x NEAA/2% B-27 (Gibco 17504-044)/90 µM 2-Merceptoethanol. For endoderm differentiation, cells were first cultured for 24h in RPMI (Gibco 21870-076)/GlutaMAX/Activin A (100 ng/ml)/Wnt3a (25 ng/ml)/0.075 mM EGTA, followed by another 48h in RPMI/GlutaMAX/Activin A (100 ng/ml)/0.2% FBS. Mesoderm differentiation was performed according to^62^, by culturing cells in basal media + 25 ng/ml Wnt3a + 50 ng/ml of Activin A on Day 1; Basal media + 25 ng/ml Wnt3a + 25 ng/ml Activin A + 20 ng/ml FGF2 on Day 2, and basal media + 25 ng/ml Wnt3a + 10 ng/ml Activin A + 20 ng/ml FGF2 + 40 ng/ml BMP4 on Day 3. Gene expression profiles were generated as described in the transcriptomic profiling section.

### High-throughput shRNA screen

Lentiviruses containing shRNA hairpins were purchased from the MISSION® shRNA Lentiviral library (Sigma) (Supplementary Table 4.1). To minimize off-target effects, a minimum of three independent hairpins were used for > 80 of candidate MRs. EpiSCs were passaged onto matrigel-coated 96-well plates (Greiner 655090) at a density of 3,000 cells/well in 100 µl CMAF medium supplemented with 10 µM Y27632. At 12h after passaging, 50 µl of the culture medium was replaced with 50 µl lentivirus with 8 µg/ml polybrene. At 24h after lentivirus transduction, the medium was replaced with 100 µl CMAF + 2 µg/ml puromycin (Sigma P7255) for selection of stable integrants; puromycin concentration was increased to 4 µg/ml at 48h. At 60h, cells were fixed with 4% paraformaldehyde (PFA) and immunostained for Oct4 (Pou5f1) (Santa Cruz sc- 5279, 1:1,000) using an Alexa Fluor 488 secondary antibody (Invitrogen A11029, 1:500). Cells were counter-stained with Hoechst 33342 (ThermoFisher H3570, 1:5,000), and plates were scanned using the IN Cell analyzer 2000 (GE Healthcare). For each well, 12 square-shaped fields were imaged for two channels, and exported images were analyzed by the IN Cell Analyzer workstation software.

### Transcriptomic profiling

To generate gene expression profiles for interactome assembly and MR analysis, cells were lysed with Trizol (ThermoFisher 15596018), and transferred into Eppendorf tubes for short-term storage at -80°C. We used 600 µl of cell lysate for total RNA extraction (Clontech 740955), followed by processing for microarray (for interactome assembly) or RNA sequencing (for differentiation signatures). For microarray analysis, total RNA was biotin-labeled using the Illumina TotalPrep RNA Amplification Kit (ThermoFisher AMIL1791) and hybridized on mouseWG-6 v2 BeadArrays (Illumina). Slides were scanned using an iScan (Illumina SY-101- 1001) to generate the resulting files.

For RNA sequencing, samples were submitted to the Columbia Genome Center for library preparation and sequencing. In brief, mRNA was enriched by poly-A pull-down, and library synthesis was performed using the Illumina TruSeq RNA prep kit (Illumina RS-122-2001). Libraries were pooled and sequenced on the Illumina HiSeq 2500 or 4000 platforms, yielding approximately 30 million single-ended reads per sample.

### PLATE-seq analyses

For PLATE-seq (Pooled Library Amplification for Transcriptome Expression), samples were prepared as described^40^, with minor modifications. Total RNA from 100 µl of Trizol-dissolved cell lysate was purified using the “No Spin” method of the MagMAX-96 for Microarrays kit (Ambion AM1830). RNA was eluted using 50 µl of elution buffer, and the concentration measured by Nanoview. 100 ng total RNA from each sample was diluted to 15 µl using nuclease-free water and transferred to 96-well plates (Applied Biosystems N8010560). We added 1 µl of 1:1,000 dilution of ERCC Ex-Fold Spike-Ins (ThermoFisher 4456739) to every other column of the plate to check for potential inter-sample contamination during library preparation.

To anneal primers for reverse transcription, 3 µl of 100 µM barcode-linked oligo-dT primer was added to each well with 5 µl of 5x ProtoScript RT buffer (New England Biolabs M0368). The plate was incubated at 94°C for 2 min and immediately cooled on ice for at least 5 min, followed by addition of the reverse transcription master mix: 2.88 mM dNTPs, 6.4 mM DTT, 0.14 U/µl SuperaseIN, 1.44 U/µl Protoscript II Reverse Transcriptase, and water to a final volume of 25 µl. The plate was then incubated at 42°C for 2h. To remove excess RT primer, 2 µl of 1:4 diluted ExoI (20 U/µl) was added to each well, and the plate incubated at 25°C for 1h. To hydrolyze the RNA, 10 µl of 1:1 mixture of 1 M sodium hydroxide and 0.5 M EDTA was added followed by incubation at 65°C for 15 min. All wells on the plate were then pooled in a 50 ml conical tube.

The pooled sample was then purified and concentrated using DNA Clean & Concentrator Kit (Zymo Research D4013) using a modified protocol. The sample was first diluted seven-fold using DNA binding buffer (Zymo Research D4003-1-L), and passed through the binding column using a vacuum apparatus. The column was washed twice with washing buffer, and eluted with 15 µl of nuclease-free water. The eluted sample was further purified using AMPure XP PCR purification beads (Beckman Coulter A63880) at a 1:1 sample to bead ratio, and eluted with 15 µl of nuclease-free water. For second-strand synthesis, 1 µl of 10 mM dNTPs and 1 µl of 100 mM adapter-linked random hexamer primer were added, followed by heating at 70°C for 2 min and immediate cooling on ice for 5 min. 2 µl of NEBuffer 2 (New England Biolabs B7002) and 1 µl of Klenow large fragment DNA polymerase (New England Biolabs M0210) were then added, followed by incubation at 25 degrees for 30 min and two rounds of purification with AMPure XP beads. The double-stranded cDNA was eluted into 15 µl nuclease-free water and concentration measured using a Qubit Fluorometer 3.0 with dsDNA high-sensitivity assay kit (ThermoFisher Q32851).

For library amplification, the cDNA was diluted to 0.05 µg/ml and PCR-amplified using Phusion DNA polymerase (New England Biolabs M0530) with 0.5 mM dNTPs, 0.5 µM Illumina RP1 and RPl1 primers. The PCR product was purified using AMPure XP beads for one round and eluted into 15 µl nuclease-free water. 2 µl of the product was analyzed by bioanalyzer with an expected broad peak from 200 bp to 1,000 bp and average length of product 500 bp. 1 nM of PCR product was hydrolyzed with an equal amount of 0.2 M sodium hydroxide, and mixed with hydrolyzed PhIX control library (Illumina FC-110-3001). The final product contained 1.8 pM library with 30% PhIX/70% sample in 1.3 ml HT1 buffer. The product was loaded on the NextSeq 500 high output v2 kit (Illumina FC-404-2005) together with two custom primers and two PhIX primers at final concentration of 300 nM each. The sample was loaded onto an Illumina NextSeq 500 sequencer for paired-end sequencing, with Read 1 (26 cycles) identifying the 8 bp barcode for sample well information and Read 2 (66 cycles) to determine the identity of the mRNA. All PLATE-seq oligo sequences are listed in Supplementary Table 10.4.

### CRISPR/Cas9 editing

To minimize off-target effects, we used a minimum of two independent sgRNAs for each candidate MR. sgRNA sequences were optimized using two online sgRNA design tools (at https://design.synthego.com/ or the site formerly available at http://crispr.mit.edu). For most experiments, we selected sgRNAs with low off-target effects, high on-target efficiency, and targeting to a common exon shared by all splice isoforms in the first half of the coding region. In some cases, we selected a pair of sgRNAs targeting intronic regions that flank a common exon located in the first half of the coding region to achieve targeted deletion of the entire exon. sgRNA primers were ordered from Integrated DNA Technologies. Single-stranded primers were annealed into double-stranded sequences with restriction enzyme engineered ends, and cloned into lentiGuide-Puro (Addgene 52963) by the Golden Gate method as described in (https://media.addgene.org/cms/filer_public/3e/e1/3ee1ce9c-99f9-4074-9a28-109d34971471/zhang-lab-sam-cloning-protocol.pdf). All inserted sgRNAs were sequence-verified.

For production of lentivirus, HEK293FT cells were grown to 90% confluency and transfected with pMD2.G, psPAX2 packaging vectors and the lentiviral expression construct, using Lipofectamine 3000 (ThermoFisher L3000001). Virus was harvested twice on two consecutive days, and titers were verified by Lenti-X GoStix Plus (Takara Bio 631280) before storage at -80°C.

We established a stable Cas9-expressing line by infecting EpiSC-5 cells with lentiCas9- Blast (Addgene 52962), followed by selection with 5 µg/ml blasticidin (Sigma 15205) for five days. Established Cas9-positive EpiSCs were confirmed for pluripotency by marker analysis as well as self-renewal and differentiation assays. To induce CRISPR/Cas9 mediated mutagenesis, Cas9-positive cells were infected with lentiGuide-Puro (Addgene 52963) inserted with sgRNA. At 24h, cells were selected with 2 µg/ml puromycin for 2 days and passaged at clonal density (1,000 cells/well on 6-well plates). At 72h after passaging, single colonies were picked into 96-well plates; at least 16 colonies were picked for each sgRNA targeted line. Colonies were grown to confluency and expanded onto 12-well plates to establish clonal cell lines. To validate targeted mutations, cells were lysed with DirectPCR Lysis Reagent (Viagen Biotech 301-C). Genomic regions were PCR-amplified (Platinum SuperFi II PCR master mix, ThermoFisher 12368050) using a pair of primers at least 200 bp away from the 5’ and 3’ ends of the sgRNA targeted site. PCR products were diluted 1:40 and 10 µl mixed with 5 µl of 5 µM sequencing primer for Sanger sequencing. Results from true clonal cell lines displayed either a single ORF consistent with identical genome editing results on both alleles, or double peaks with equal height indicating different alleles on the homologous chromosomes. In the latter case, to determine sequences of the individual alleles, PCR products were cloned into TOPO vector (Strataclone Blunt PCR Cloning Kit, Agilent 240207) and selected on agar plates with ampicillin and kanamycin. Multiple colonies were mini-prepped (Qiagen 51306) and sequenced.

### RT-qPCR

Total RNA was reverse-transcribed to cDNA using SuperScript Reverse Transcriptase II. Diluted cDNA was mixed with SYBR green master mix (Applied Biosystems A25777) and qPCR reaction was performed on an ABI7500 instrument (Applied Biosystems) using conditions of 50°C for 2 min; 94°C for 2 min; 94°C for 30 s + 60°C 30 s for 40 cycles. The expression of *Gapdh* or *Hprt* was used to normalize the loading error, and the Delta-delta Ct method was used to quantify relative gene expression levels.

### Immunofluorescence and immunohistochemistry

For adherent cells, cells were fixed with 4% PFA for 15 min and washed with 1x PBS. Cells were permeabilized by 0.1% TritonX-100 for 10 min. Blocking was performed using 5% goat serum or 2% non-fat milk, and primary antibody was diluted with 1% serum or non-fat milk for staining overnight at 4°C. Cells were then washed three times with 0.05% Tween-20 and secondary antibody was diluted 1:500 with 1% goat serum or non-fat milk for staining at room temperature for 1 hour, followed by washing three times with 0.05% Tween-20. Nuclei were stained with Hoechst 33342 (1:5,000 dilution in PBS) for 10 min and washed with PBS twice.

For teratomas, hematoxylin and eosin (H&E) staining or immunohistochemistry was performed on 5-micron paraffin sections. For H&E staining, sections were rehydrated by serial incubation with xylenes three times, 100% EtOH twice, 95% EtOH twice, 70% EtOH once, and 50% EtOH once. Slides were dipped into hematoxylin for 3 min, followed by rinsing with deionized water and quick incubation with acid ethanol. Slides were then incubated with eosin for 30 s and then dehydrated by incubating with solvents in the reverse order of rehydration. The slides were then cover-slipped using Clearmount.

For immunohistochemistry, rehydrated slides were blocked with 3% hydrogen peroxide for 20 min. Antigen retrieval was performed by heating in a steamer for 45 min. Slides were then permeabilized with PBS + 0.1% Triton-X for 15 min and blocked with 5% serum for 1 hr. Primary antibody was diluted with 1% serum and incubated with slides in humid chamber overnight at 4°C. Secondary antibody staining used biotinylated antibodies at 1:500 dilution in 1% serum. Color detection used an ABC amplification kit and Vector NovaRed Substrate Kit (Vector Laboratories). Hematoxylin staining was performed by quick dipping slides into hematoxylin 3-5 times, followed by washing, dehydrating, and coverslipping as described above.

### Western blotting

Attached cells were grown to confluency on 10 cm^2^ dishes and dissociated with 0.05% trypsin. Cell suspension was lysed, measured and denatured by following protocol as described in PMID: 29625057. For SDS-PAGE, a 10-20 µg sample was loaded onto Bio-Rad pre-cast 4%-20% Mini-PROTEAN TGX gels (4561094S), run for 2-3 hours at 120-140V, and then transferred to PVDF membranes at 400 mM for 3 hours. Primary antibody staining was performed overnight at 4°C with antibodies diluted in 5% non-fat milk in TBST. HRP-conjugated secondary antibody was used to incubate the membrane for 1 hour at room temperature. Color reaction was performed using SuperSignal West Pico Plus Chemiluminescent Substrate (ThermoFisher 34577).

### Colony formation assays

Confluent cells were dissociated, counted (Bio-Rad TC20), and replated at clonal density (500 cells/well) on 12-well attachment plates, in 1:2 diluted CMAF without Y27632. Media was refreshed once at 48h after replating. At 96h, cells were stained with methylene blue by incubating with 1% methylene blue diluted in 50% methanol for 5 min followed by washing with tap water for 10 min. Alternatively, cells were stained for alkaline phosphatase using a commercially available kit (Stemgent 00-0055). In a separate experiment at 120h after plating at clonal density, cells were dissociated with 0.05% trypsin and resuspended in 1 ml basal media. Cell numbers in suspension were counted by the Bio-Rad TC10 cell counter and divided by the initial number of plated cells to determine the fold-change.

### Embryoid body formation

Embryoid body formation was performed as described^63–65^, with minor modifications. In brief, EpiSCs were dissociated into single cells and diluted to 10,000 cells/ml in EB growth media containing DMEM/F12, 20% FBS, 1x MEM non-essential amino acids, 1x GlutaMAX, and 55 µM 2-merceptoethanol. Hanging drops were plated as 30 µl drops containing 300 cells on the top of a 150 mm culture dish. After 72h, embryoid bodies were washed and imaged.

### *In vitro* lineage-specific differentiation analysis

EpiSCs were differentiated into neuroectoderm and mesendoderm lineages as described^66,67^, with minor modifications. In brief, 500-5,000 cells, depending on their growth rate, were plated in 24-well culture plates with 0.5 ml CMAF + 10 µM Y27632. The next day, differentiation was initiated in MEF conditioned media, lacking FGF2 and Activin A. For neuroectoderm differentiation, cells were treated with 10 µM SB431542 (Millipore Sigma S4317) + 12 ng/ml FGF2 for 7 days. For mesendoderm differentiation, cells were treated with 10 µM SU5402 (Millipore Sigma 572630) + 5 ng/ml Activin for 3 days, followed by 10 ng/ml BMP4 + 20 ng/ml FGF2 + 30 ng/ml Activin for 4 days. On day 4, after most wells reached confluency, cells were dissociated and replated onto 24-well and 96-well plates to achieve 30-50% confluency. Cells continued differentiation for another three days before harvesting for RT-qPCR analysis (24- well cultures) and immunofluorescence staining (96-well cultures).

### Image analysis

Immunofluorescence-stained samples in 96-well plates (Greiner 655090) were imaged by the Cytation 5 Cell Imaging Multimode Reader using the Biotek Gen5^TM^ software. For each marker, exposure time for two channels (GFP: 488 nm and DAPI: 405 nm) was adjusted to reduce background and to maximize on-target signal. For each well, 9 square-shaped field were recorded for each channel and exported to a stitched image file. Two channel images from the same sample were pseudo-colored and overlaid by Fiji ImageJ.

An image analysis pipeline was developed using the CellProfiler software (version 3.0.0). This pipeline scans the DAPI-channel image to mark the area containing the cell nucleus, and then scans the 488nm-channel image to quantify the nuclear staining intensity of the marker protein. For markers that stain cytoplasm such as Doublecortin, the area surrounding the nucleus is marked for green fluorescent intensity quantification. CSV formatted files containing green fluorescent intensity quantification of individual cells in each sample well were exported, which were then analyzed by an R pipeline generated by the Herbert Irving Comprehensive Cancer Center (HICCC) Confocal and Specialized Microscopy Shared Resource. In general, the pipeline firstly determines the background staining intensity for each antibody marker, and then reads through the CSV file and identifies cells that show higher intensity of green fluorescence signal than the background. The proportion of cells with positive marker expression is then calculated by dividing the number of cells with higher than background staining to the total number of cells recorded. The proportions, expressed as percentages, are then normalized to the batch-matched control samples (staining on wild type EpiSCs). Comparing biological replicates of mutants to controls determines the statistical significance (Student’s T-test, two-tails, unpaired).

### Teratoma formation

Teratoma formation was performed as described^68^ with minor modifications. In brief, EpiSCs were dissociated into single cells, and 300,000 cells were resuspended in 50 µl Matrigel. Cells were injected into the gastrocnemius muscle of immunodeficient mice, and mice were monitored daily to track tumor growth. Mice were euthanized when tumors grew to 2 cm diameter, around 25 days after injection. Tumors were dissected intact, weighed, cut into 1 mm sections by razor blade, fixed with 4% PFA or 10% formalin, paraffin-embedded, and processed for H&E staining or immunohistochemistry. All animal studies were approved by and conducted according to standards set by the Columbia University Institutional Animal Care and Use Committee (IACUC).

### EpiSC perturbation (pilot project)

To guide the design of the EpiSC perturbation assays for interactome generation, we performed a pilot experiment. Specifically, Illumina BeadArrays (mouse WG-6, Illumina) were used to profile the transcriptome of 33 independent EpiSC samples. This experiment was divided in three arms. The first arm consisted of cells perturbed with 10 small molecule compounds (100 nM rapamycin, 250 nM PF573228, 2 μM dorsomorphin dihydrochloride, 25 μM BAY 11-7082, 50 μM forskolin, 5 μM BAPTA-AM, 500 pM docetaxel, 50 nM wortmannin, 2 μM purmorphamine, and 2 μM SU5402) and vehicle control for 24h. For the second arm, cells were first perturbed for 36h with 5 μM RA and for 24h with 10 small molecule compounds and vehicle control. Specifically, the cells were exposed to either control media or RA for 12h, small molecule compounds were then added, and cells were further incubated for 24h. The third arm consisted of the same RA/compounds combination experiment but was performed using an inverted perturbagen vs. RA treatment order (*i.e.*, 24h RA and 36h perturbagen treatment). Gene expression profiles were generated using an iScan BeadArray scanner (Illumina) and pre-processed by variance stabilization and robust spline normalization, as implemented by the lumi R package^69^. Gene expression profiles are available from the Gene Expression Omnibus (GSE197632).

A 2D plot (Extended Data Fig. 1a)—obtained by projecting the data on the first two principal components, after scaling by subtracting the mean and dividing by the standard deviation of each gene—showed that treating first with perturbagens and then with RA induced greater data dynamic range than the opposite order. We thus decided to perform the larger study by first treating with each perturbagen for a total of 36h, and with each morphogen for 24h, thus maximizing the dynamic range for the reverse engineering of an EpiSC-specific interactome (see below).

### Generating an EpiSC context-specific interactome

The transcriptomes of the two EpiSC cell lines—following treatment with combinations of 33 perturbagen (36h) and 5 morphogens (24h), in two growth media conditions (CMAF and Basal)—were profiled by Illumina BeadArrays (mouse WG-6 v2, Illumina). Poor quality profiles were removed, for a total of 276 high-quality gene expression profiles. (Supplementary Table 2.1). Gene expression profiles were generated using an iScan BeadArray scanner (Illumina) and pre- processed by variance stabilization and robust spline normalization, as implemented in the lumi R package^69^. Gene expression profiles are available from the Gene Expression Omnibus (GSE197414).

The profiles were then analyzed using the ARACNe algorithm^24,70^ to reverse engineer an EpiSC-specific transcriptional interactome representing the transcriptional targets of 1,478 regulatory proteins. The latter included those annotated as GO:0003700 (*transcription factor activity*), GO:0004677 (*DNA binding*), and GO:0030528 (*transcription regulator activity*), or as GO:0004677 and GO:0045449 (*regulation of transcription*) in the Gene Ontology (GO) Molecular Function database^37^. To generate an optimal consensus network, we performed 100 bootstrap iterations of the ARACNe analysis. Parameters were set to DPI = 0 (data processing inequality tolerance) and MI P-value *p* ≤ 10^−8^ (mutual information threshold). The resulting interactome comprises 911,753 transcriptional interactions (Supplementary Table 3.1).

### Predicting master regulators of pluripotency and differentiation

To generate time courses representing the path from a pluripotent to a differentiated cell state, we generated RNA-Seq profiles of EpiSC-5 cells undergoing 5 distinct differentiation treatments (differentiation arms), at 3h, 6h, 12h, 18h, 24h, 36h, 48h and 72h (Supplementary Table 2.2). RNA-Seq profiles were generated with the Illumina TruSeq protocol, using a HiSeq 2000 instrument (Illumina), at the Columbia Sulzberger Genome Center. Sequencing data were mapped to the mouse genome MGSCv37 with TopHat v1.3.3^71^. Reads mapping to known genes, based on Entrez gene identifiers, were then counted using the GenomicFeatures R package, from Bioconductor^72^. Summarized raw counts for 23,283 known genes in 144 samples are available from the Gene Expression Omnibus (GSE199114). We computed a size factor for each sample using the median log-ratio method and identified as outliers the samples showing residuals from a linear fit of the computed size factors vs. sequencing depth (total number of mapped reads) greater than 50%. Samples identified as outliers were removed from further analyses. Gene expression data were then normalized by equivariance transformation, based on the negative binomial distribution, with the DESeq R package^73^.

The expression likelihood for each gene was estimated by fitting a mixture of three gaussian distributions to the Log (RPKM + 1) data, across all genes in the integrated dataset, i.e. raw counts for each gene aggregates across all control samples, with the mixtools package ^74^, and expressed as 1 minus the relative likelihood for an observation, expressed as Log (RPKM + 1), to be derived from the first (lowest mean) distribution.

For each differentiation arm, gene expression signatures were computed at each time point by differential gene expression analysis against the profiles of undifferentiated EpiSC-5 cells (t = 0h) as the reference, using Student’s t-test (Supplementary Table 4.3). Protein activity signatures were then assessed using the msviper function of the VIPER R package, available from Bioconductor^25^. Normalized Enrichment Score (NES) and associated *p*-values were estimated by permuting sample labels uniformly, at random 1,000 times. For time points with fewer than 5 replicates, the closest samples from contiguous time points were included, up to 5 sample in total, to support the permutation-based estimation of NES and *p*-values (Supplementary Table 4.4). These were assessed based on the similarity of their gene expression signatures using Euclidean distance.

For each differentiation arm and time point, differential protein activity *p*-values were assessed from the VIPER-computed NES. To account for proteins that could be activated in some conditions and inactivated in others, *p*-values were further integrated using the Fisher’s method. Finally, to compensate for a potentially non-statistically independent contribution of the mesoderm and endoderm differentiation arms—which produced strongly correlated protein activity signatures—we down-weighted their contribution by half during Fisher’s p-value integration.

The resulting ranked protein list identified 300 candidate MRs (p < 10^-4^) (Supplementary Table 4.1). Among these, 134 (45%) had been previously implicated in pluripotency regulation in the literature (Supplementary Table 4.2). To prioritize candidate MRs for an experimentally feasible shRNA-based validation assay, we selected candidate genes using the following four criteria: (1) The top 120 most differentially active proteins, as ranked by p-value, were included by default; (2) 70 additional proteins were selected uniformly at random, from proteins ranked between positions 121 and 300 by p-value; (3) 30 previously established pluripotency regulators— including 18 that were not identified among the top 300 MRs, and 12 that were not represented as transcriptional regulators in the EpiSC interactome—were also included; (4) Finally, since MR validation was based on silencing assays, we removed those not expressed in EpiSC (likelihood > 0.5). Taken together, these criteria produced a list of 172 candidate MRs for experimental validation (Supplementary Table 6.2). Note that of the 18 previously identified pluripotency regulators that were not in the original list of 300 candidate MRs, only 4 (Gli2, Jarid2, Sall4, and Sox2) remain among the 132 MRs used in the final network analysis. Thus, 128 out of the 132 MRs in the final network were identified in a completely unbiased manner.

### Dimensionality reduction analysis

To project the data in 2 dimensions, we used Principal Component Analysis (PCA), as implemented by the *prcomp* function of the stats package in R-system. We also used the ‘umap’ v0.2.7.0 R package to perform Uniform Manifold Approximation and Projection (UMAP) analysis, as available from the CRAN repository (https://cran.r-project.org/web/packages/umap/index.html).

### Oct4 immunofluorescence data analysis

We used the IN Cell Image Analyzer to perform immunofluorescence intensity quantification at the individual cell level. This instrument can scan wells in a 96-well plate and exports image files using two channels (Green Fluorescence (GF) and DAPI). These data were analyzed using the IN Cell 2000 workstation software. By setting up diameter parameter as 5um∼10um, the software first analyzes the DAPI-channel to identify individual nuclei. The green fluorescence of each nucleus is then quantified by scanning the 488 nm channel. CSV files comprising the green fluorescence quantification for the cells in each well were then exported by the software. The data was acquired in 22 96-well plates (batches), with each plate assaying a subset of MR-directed and negative control shRNA hairpins.

Overall, the instrument quantified the fluorescence of 23,317,327 individual cells. These include 6,537,819 cells representing control conditions, 15,243,283 cells transduced with candidate MR-targeting shRNA hairpins, and 1,536,225 cells transduced with shRNA clones targeting established pluripotency regulators. In total, 623 and 47 hairpins, targeting 158 candidate MR and 14 established pluripotency regulators, were tested, respectively.

Cells with fluorescence values higher than 6 inter-quartile range (IQR) units above the median of control conditions were removed from further analysis as outliers. The data were then normalized, independently for each plate, using the plate-specific control conditions. A cumulative empirical density distribution was computed for the control conditions of each plate and used to estimate the normalized fluorescence z-score of each cell. By using plate-specific controls, the analysis eliminated all systematic difference between plates (batch effect). Data were summarized on an individual well basis by computing the area over the cumulative empirical distribution across all cells in the well, thus eliminating well-specific cell density as a confounding factor. Next, the statistical significance for the differential Oct4 protein level was estimated by comparing the summarized replicates for each hairpin to the summarized replicates of control wells, using Student’s t-test. Finally, to compute a gene-level z-score, the statistics of each associated shRNA clone were integrated using Stouffer’s method.

To account for different silencing efficiencies, the shRNA-mediated reduction in the activity of the target protein (see next section below), expressed as z-score, was used to weight the contribution of the corresponding shRNA. Thus, shRNAs failing to inhibit the target proteins provided no contribution to the score. The final Oct4 *corroboration score* (CS) was computed by multiplying the Stouffer’s integrated z-scores for the corresponding gene by the sign of the MR protein activity change during differentiation (i.e., +1 for activity increase and -1 for activity decrease). Finally, we derived the *Oct4 protein score* as the log-transformation of CS while maintaining its sign:

Oct4 protein score = log_2_(|CS| + 1) × sign(CS)

### EpiSC expression profiles after candidate MR silencing

We used the PLATE-seq technology^40^ to generate gene expression profiles from EpiSC cells, at 72h following lentiviral-mediated shRNA transduction. Sequencing reads were de- multiplexed as previously described^40^, mapped to the mouse genome build GRCm38 (mm10), and counted at the known gene level with STAR aligner v2.5.2 ^75^. Summarized gene expression counts are available from the Gene Expression Omnibus database (GSE199855). Expression data were normalized by equivariance transformation, based on the negative binomial distribution with the DESeq R-system package^73^.

For each candidate MR-targeting shRNA clone, differential gene expression signatures were computed, against plate-matched, non-targeting controls, as a reference. Statistical significance was assessed by moderated Student’s t-test, as implemented in the limma package from Bioconductor^75^ and represented as a z-score vector.

For each MR-targeting shRNA, differential protein activity was assessed by VIPER analysis, using the EpiSC interactome. To control for differences in regulon size (number of transcriptional targets per regulon), all regulons were capped to the top 50 targets, based on their interaction likelihood.

### Inverted regulon analysis

We used the silencing assays to identify regulons likely to predict an inverted activity for the associated regulatory protein. Specifically, we assessed the change in VIPER-assessed protein activity following MR silencing and changed the regulon sign of candidate MRs, whose activity was assessed as increased following silencing. This addresses a problem originally reported in the VIPER manuscript^25^, whereby the activity change may be inferred in the opposite direction due to negative autoregulatory loops that may anticorrelate the expression of a regulator and that of its targets. To identify the regulons reporting an inverted activity, we fitted a mixture of two gaussian distributions to the empirical distribution of estimated activity z-scores, for all silenced regulators, representing the distribution of differential protein activity for the inverted and non-inverted regulons. Then, based on these two gaussians, we estimated a conservative threshold to identify regulons with an inverted regulatory sign. This was computed as the z-score corresponding to a likelihood ratio of 4 between the gaussian models fitted to the inverted and non-inverted regulons, respectively.

### Differentiation signature score

We then assessed whether the changes induced by candidate MR silencing would recapitulate those observed in the differentiation time-courses. This was achieved by computing the enrichment of the 100 most differentially active transcriptional regulators, following shRNA-mediated silencing, in proteins differentially active at 72h in the morphogen-mediated differentiation assays, using the aREA algorithm^25^. For these analyses, protein activity was again assessed using regulons capped at the top 50 transcriptional targets.

The Normalized Enrichment Score (NES) of each MR-specific hairpin and for the 5 lineage specific differentiation assays, was integrated using a weighted implementation of Stouffer’s method. Mesoderm and endoderm differentiation were included with half the weight of other lineages since they were strongly correlated. In addition, as discussed above, the efficiency of each hairpin in reducing the activity of the target protein, expressed as z-score, was also used to weigh its contribution. The final NES was multiplied by the sign of the MR protein activity change, change to obtain the *differentiation z-score* (DZ), such that concordant and discordant effects would have a positive and negative NES, respectively. Finally, we derived the *Differentiation Score* as the log-transformation of DZ while maintaining its sign:

Differentiation Score = log_2_(|DZ| + 1) × sign(DZ)

### MR interactome refinement

To refine the transcriptional targets of each candidate MRs using experimental data, we integrated z-scores corresponding to (1) the differential expression (DE) of its candidate targets, following shRNA-mediated silencing, and (2) the mutual information (MI) between the MR and its candidate targets (see EpiSC context-specific interactome above). To avoid bias in assessing the statistical significance of these metrics, we rank-transformed and scaled between 0 and 1 both variables before integration, while preserving the differential expression (DE) score sign. Integration between MI and DE was performed using a weighed implementation of the Stouffer’s method. The DE contribution was weighted by the silencing efficiency of each shRNA hairpin, as previously discussed. Such weights were computed by a logistic transformation of the silencing efficiency scores with inflection point equal 1 and trend at the inflection point equal 3. MI scores were only limited to the regulator’s transcriptional interactions present in the original EpiSC interactome, in this way making full use of the data processing inequality filter. The “TFmode” component of the new regulons^25^ was computed for each target as the average between the direction-corrected “TFmode” from the EpiSC interactome and the DE score for the target in response to the regulator silencing.

To estimate the “likelihood” component of the new regulons, we first computed two integration scores: (1) a non-directional integration score (NDIS) as the weighted Stouffer’s integration of the rank-transformed MI and |DE|, where |x| denotes the absolute value of x, and (2) a directional integration score (DIS) as the weighted Stouffer’s integration of the rank-transformed MI multiplied by the sign of the direction-corrected “TFmode” from the EpiSC interactome and the DE score. Finally, the direction-corrected “TFmode” from the EpiSC interactome was used to weight the contribution of the NDIS and DIS scores when computing the integrated interaction score (IS) as DIS x |TFmode| + NDIS x (1 - |TFmode|). The “likelihood” component was then estimated by scaling the IS by its maximum value across all the targets for the regulator. The top 200 transcriptional targets per regulator, based on IS, were included in the interactome. For VIPER analysis, we limited the number of transcriptional targets per regulon such as the effective size— *i.e.*, sum(likelihood^2)—was capped at 50.

### Selection and evaluation of pluripotency MRs

In the first assay using Oct4 protein level as the readout for pluripotency alteration following silencing, we tested 172 selected genes expressed in EpiSCs, including the top 95 most differentially active candidate MRs expressed in EpiSC, 63 additional candidate MRs expressed in EpiSC and ranking between 121 and 300, and 14 additional genes previously known to be involved in pluripotency. An additional candidate MR, Kmt2a, could not be evaluated because all hairpins used for targeting did not pass the data QC in either of the two assays below. From the 172 genes tested, 117 MRs (68%) were found to down-regulate Oct4 protein levels (FDR < 0.05) and 15 MRs (8.7%) were found to up-regulate Oct4 when silenced. In a second, parallel assay using PLATE-seq datasets from silenced cells, we obtained analyzable datasets after QC for 154 of the 172 MRs—including all the top 95 most differentially active candidate MRs expressed in EpiSC, 54 candidate MRs ranking between 121 and 300, and 5 genes previously known to be involved in pluripotency—following their individual silencing in EpiSCs. Among those 154 MRs, a total of 132 were confirmed by PLATE-seq based assay to induce an expected differentiation state when silenced (positive DS with FDR < 0.05; Fig. 2d).

### Experimentally refined network analysis

Based on the new experimentally-refined network, we used VIPER to assess the differential activity of 132 candidate pluripotency MRs following shRNA-mediated silencing. The NES scores for each shRNA targeting the same candidate MR were integrated using the Stouffer’s method, weighted to account for the hairpin silencing efficiency. A major advantage of this approach is that we could determine network directionality. ARACNe cannot determine the directionality of an interaction between two regulatory proteins A and B. However, if B’s expression or activity is affected by the experimental silencing of A, or vice-versa, this can effectively resolve this issue, thus leading to a bona fide *causal network*. This can also resolve situations where A regulates B and B regulates A (autoregulatory loop).

The statistical threshold to determine the presence of a causal edge was determined by evaluating the relationship between the number of nodes (proteins) and edges at different thresholds. Based on this analysis, we inferred an edge between a silenced regulator P*_r_* and a downstream target protein P*_t_* conservatively, only when the activity of P*_t_* significantly changed in response to P*_r_* silencing (FDR < 10^-10^). The final network comprises 120 MRs and 1,273 causal MR→MR edges (Fig. 4a). Of the validated MRs, 12 were not included in the network because no significant interactions with other MRs were identified by the analysis.

### Differentiation phenotype analysis

As a secondary validation assay, aimed at assessing whether a differentiation phenotype would emerge following CRISPR/Cas9 based silencing, we focused on 70 of the 132 MRs that had not been previously reported as pluripotency MRs in the literature, as well as 4 additional MRs that were previously known. A total of 1,150 clonal lines were generated by sgRNA targeting each one of these MRs, together with 128 scramble/mock controls. Among them, 520 lines contained biallelic frameshift mutations. Analyzing the morphology of those 520 lines identified 15 MRs whose knock-out significantly affected EpiSC morphology (Supplementary Table 7.1). In addition, we could not recover biallelic frameshift lines for 5 additional MRs (Chi-square test, p < 0.05; Extended Data Fig. 2a), suggesting that null mutants for these genes may be impaired in self-renewal and/or clonogenicity. Collectively, a total of 15+5 = 20 MRs were validated by CRISPR/Cas9 based analysis, including 3 of 4 genes selected as known MRs (75%) and 17 of 70 novel MRs (24%). As a follow-up, a panel of *in vitro* functional assays evaluating self-renewal and differentiation potential of these 20 MRs were performed using 40 knock-out (KO) lines for the 15 MRs with biallelic frameshifts and 11 knock-down (KD) lines for the remaining 5 MRs, as well as 9 empty or mock controls (Supplementary Table 7.2, 7.3). 2 MRs (*Cbl* and *Zc3h13*) display the strongest results and their KO lines were further evaluated by *in vivo* teratoma formation assay.

### Causal pluripotency MR network analysis

Community analysis was performed with the Louvain multi-level modularity optimization algorithm ^50^, as implemented in the igraph package in Bioconductor. For each node, we computed the betweenness centrality, defined as the number of shortest paths connecting all node pairs in the network that pass though the evaluated node; the degree, as the number of edges associated to each node; the in-degree, as the number of incoming edges for each node; and the out-degree, as the number of outgoing edges for each node, using the implementation for computing these centrality measurements in the igraph package. We defined the outlier nodes showing the highest betweenness as “Mediators”, because they mediate most of information flow in the network.

We define the Regularized out-degree (ROD) for each node as the total number of outgoing interactions (out-degree) multiplied by the difference between its out- and in-degree, after scaling such differences between 0 and 1 along all nodes. The ROD analysis identified three proteins classes including (a) Speakers, defined as nodes with high levels of ROD, (b) Communicators, defined as nodes with intermediate ROD, and (c) Listeners, defined as nodes with low ROD. The ROD thresholds for considering a node as a Speaker, Communicator or Listener were inferred by fitting a mixture of 3 gaussian distributions to the ROD distribution along all nodes. We defined the ROD threshold separating listener from communicator nodes as the point of equal probability between the first (lowest mean) and second fitted distributions. The speakers were defined by showing an ROD higher than the quantile of the third fitted distribution corresponding to a right-tail probability lower than 5% (Fig. 5b).

### Community functional gene-set analysis

The enrichment of the MR silencing gene expression signatures on Gene Ontology Biological Processes (GO-BP), or on MsigDB v7.4, including KEGG, Reactome, Biocarta, PID pathways and biological hallmarks were estimated with the aREA algorithm. Enrichment due to gene-sets overlap was corrected using shadow analysis as implemented in the VIPER package from Bioconductor^25^. Multiple hypothesis testing was addressed by False Discovery Rate (FDR). The enrichment results were integrated for each community, or each category of regulator (i.e. speaker, communicator and listener) using average integration of the normalized enrichment scores weighted by the community score. The results are reported for all GO-BP or MsigDB categories significant in at least one community, or at least one speaker category at FDR < 0.01. Due to the low number of hits obtained from MsigDB dataset, only the enrichment of GO-BP to the four communities were reported in Supplementary Table 9.1. The GO-BP terms preferentially enriched to each of the four communities were hand-picked using the following criteria: 1) GOBP terms with -log_10_ p < 2.5—a value selected to optimize the number of statistically significant categories—in all four community signatures were first filtered out; 2) For the remaining GOBP terms, the maximal number of the -log_10_ p value among all four communities were determined, and the differences between this maximal value to the other three values were calculated; 3) GOBP terms with maximal value larger than any other three values by more than 2, or GOBP terms that are significantly enriched to only one community (-log_10_ p < 2.5) but not the other three communities were included in Supplementary Table 9.2 and Fig. 4h.

### Dataset availability

The raw and normalized expression datasets are available from Gene Expression Omnibus: (1) EpiSC perturbed with retinoic acid and small molecule compounds at 24h and 36h (n=33, GSE197632); (2) EpiSC perturbed with 5 morphogens and small molecule compounds used for EpiSC interactome reverse engineering (n=276, GSE197414); (c) Differentiation time course for EpiSC (n=144, GSE199114); (d) shRNA-mediated targeting of 154 genes in EpiSC (n=798, GSE199855). The data for the immunofluorescence quantification of Oct4 protein levels after shRNA-mediated silencing of 172 genes is available in github (https://github.com/reef103/manuscript-episc).

### Code availability

All the code used for analysis and graphical representation, as well as the required data is available from github (https://github.com/reef103/manuscript-episc).

## Extended Data Figure Legends

**Extended Data Figure 1:**
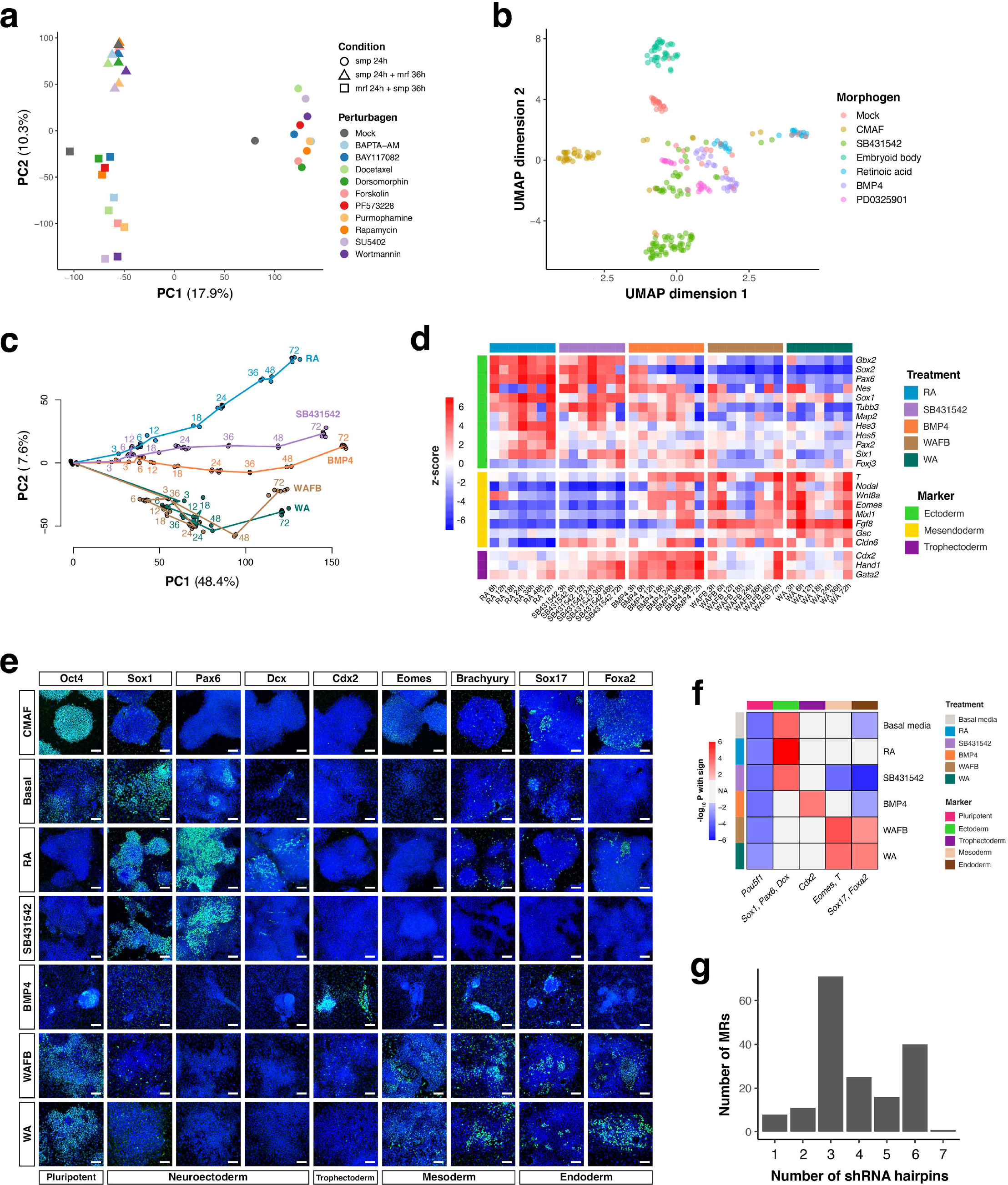
Interactome assembly, master regulator prediction, and shRNA- based validation. (a) Principal components analysis (PCA) for expression profiles of EpiSCs perturbed with the indicated small-molecule compounds, morphogens, and combinations. The proportion of variance captured by each component is indicated. (b) Uniform Manifold Approximation and Projection (UMAP) projection of 276 EpiSC expression profiles obtained after treatment with different combinations of morphogens and small-molecule perturbagens. (c) PCA of the time course expression profile data for five differentiation treatments. Numbers next to the points represent hours after the treatments. (d) Heatmap showing differential expression of 23 lineage-specific markers during the differentiation time course. Differential expression is shown as z-scores relative to 0 hr EpiSC untreated controls. (e) Immunofluorescence staining of marker expression at 72h of differentiation, as well as untreated (CMAF) and basal media controls. Marker expression is shown in green and DAPI staining in blue. Scale bars: 100 microns. (f) Heatmap showing down-regulation (blue) and up-regulation (red) of marker expression for each treatment in panel e to untreated (CMAF) controls (-log_10_(FDR), 2-tailed U-test). Statistical significance for a panel of markers was assessed by integrating individual z-scores with Stouffer’s method; only statistically significant scores (FDR< 0.05) are colored. (g) Number of independent shRNA hairpins used to target each MR (n = 172) in PLATE-Seq and Oct4-INCell assays.

**Extended Data Figure 2:**
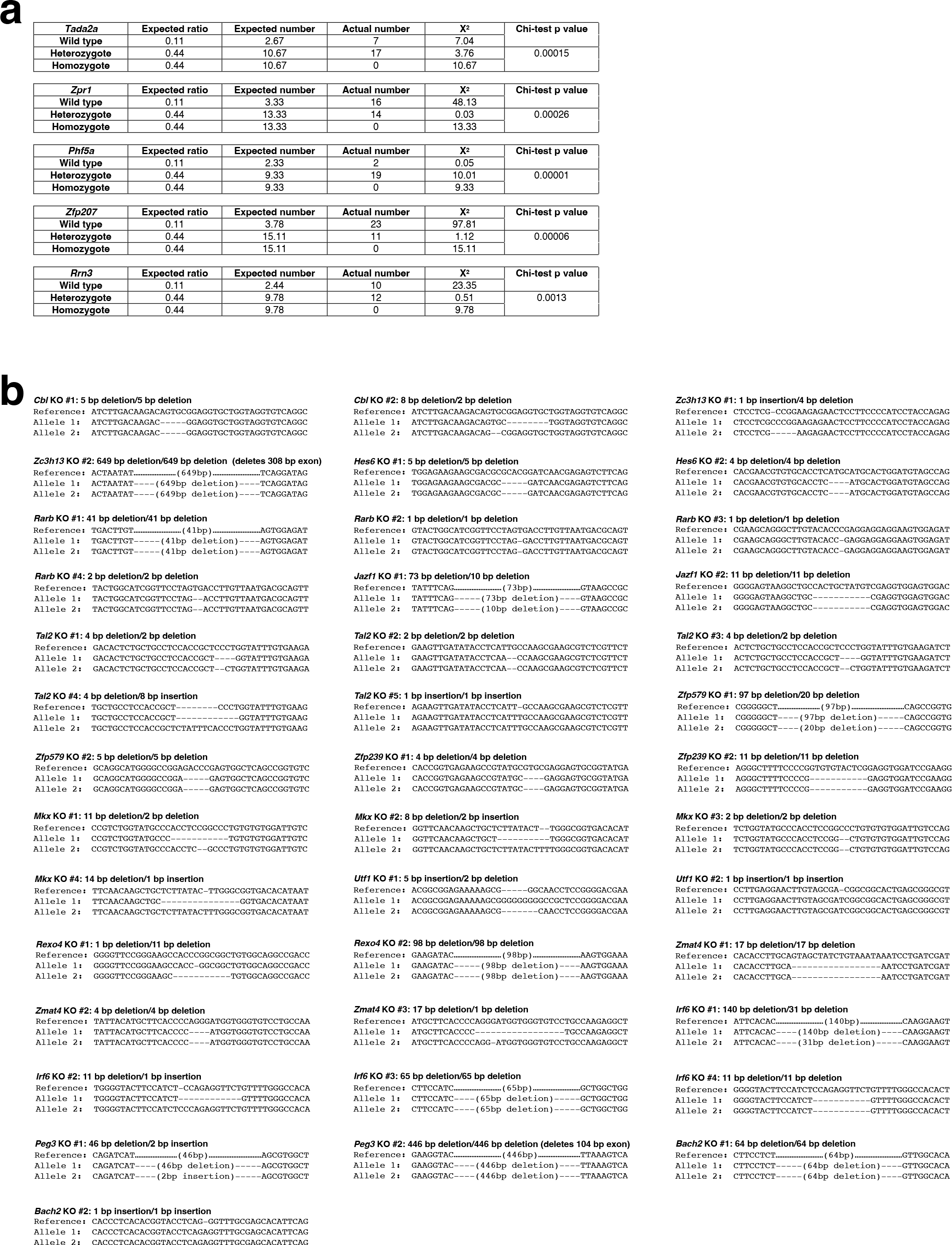
Genotyping of CRISPR KO EpiSC lines. (a) Chi-squared test results obtained by comparing the observed to the expected frequency of each genotype in the cell lines targeted for Tada2a, Zpr1, Phf5a, Zfp207 and Rrn3. P values represent the integrated statistical significance of the X^2^ values of heterozygote and homozygote groups (df = 1). (b) Genomic sequence of the sgRNA-targeted locus in the KO cell lines after CRISPR/Cas9-mediated targeting. Results for 40 targeted lines selected for functional characterization are displayed.

**Extended Data Figure 3:**
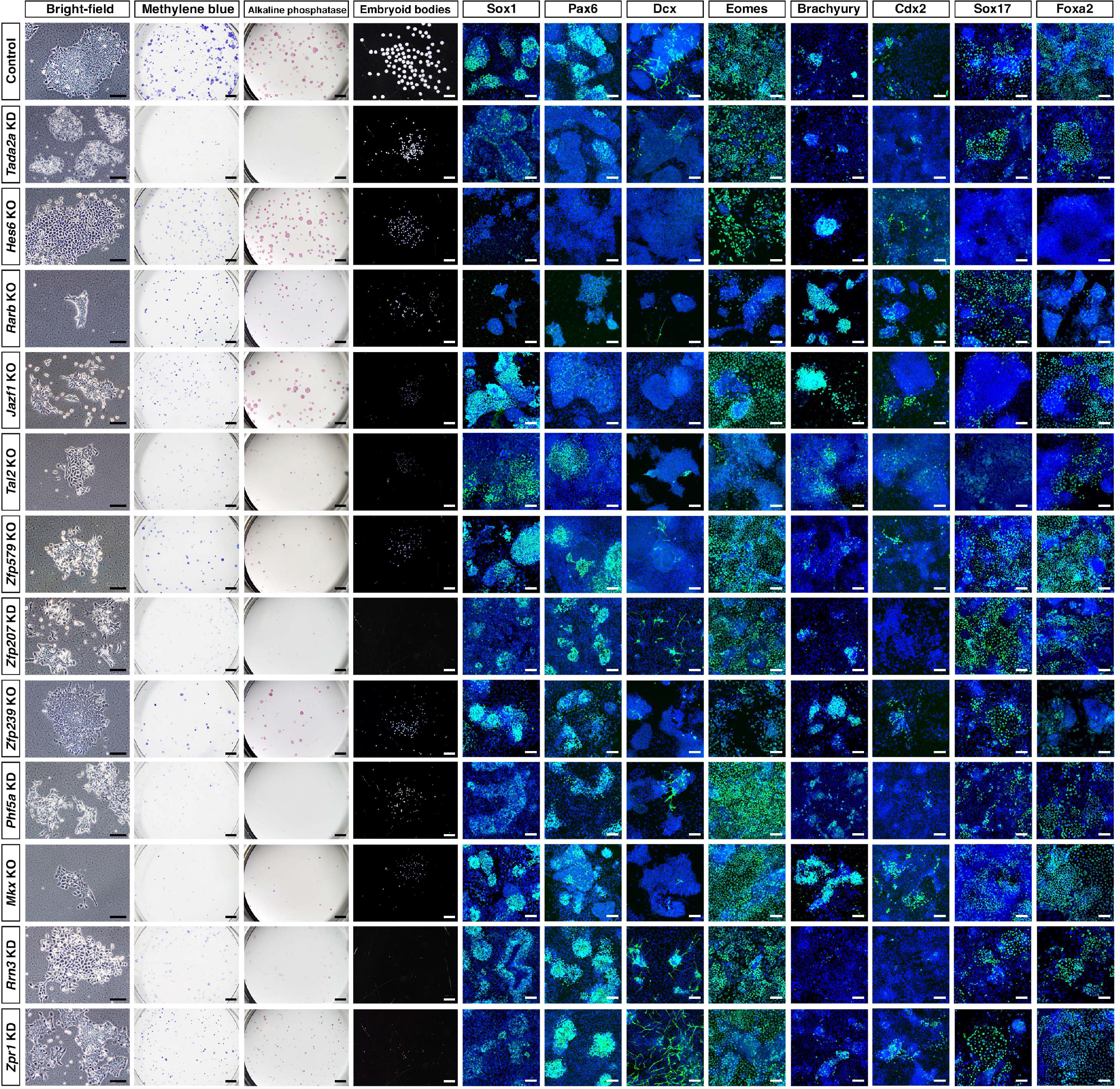
Functional analysis of master regulators (part 1). Analysis of a control EpiSC line and 12 CRISPR KO or shRNA-silenced (KD) lines in self-renewal and differentiation assays. Shown are morphology in pluripotency-maintaining culture condition (bright field); colony formation assayed by methylene blue and alkaline phosphatase staining; embryoid body formation after 72h of hanging drop culture; expression of neuronal lineage markers at 7 days after induction of neuroectoderm differentiation conditions; expression of mesoderm and endoderm lineage markers at 7 days after induction of mesendoderm differentiation. Scale bars: 100 microns.

**Extended Data Figure 4:**
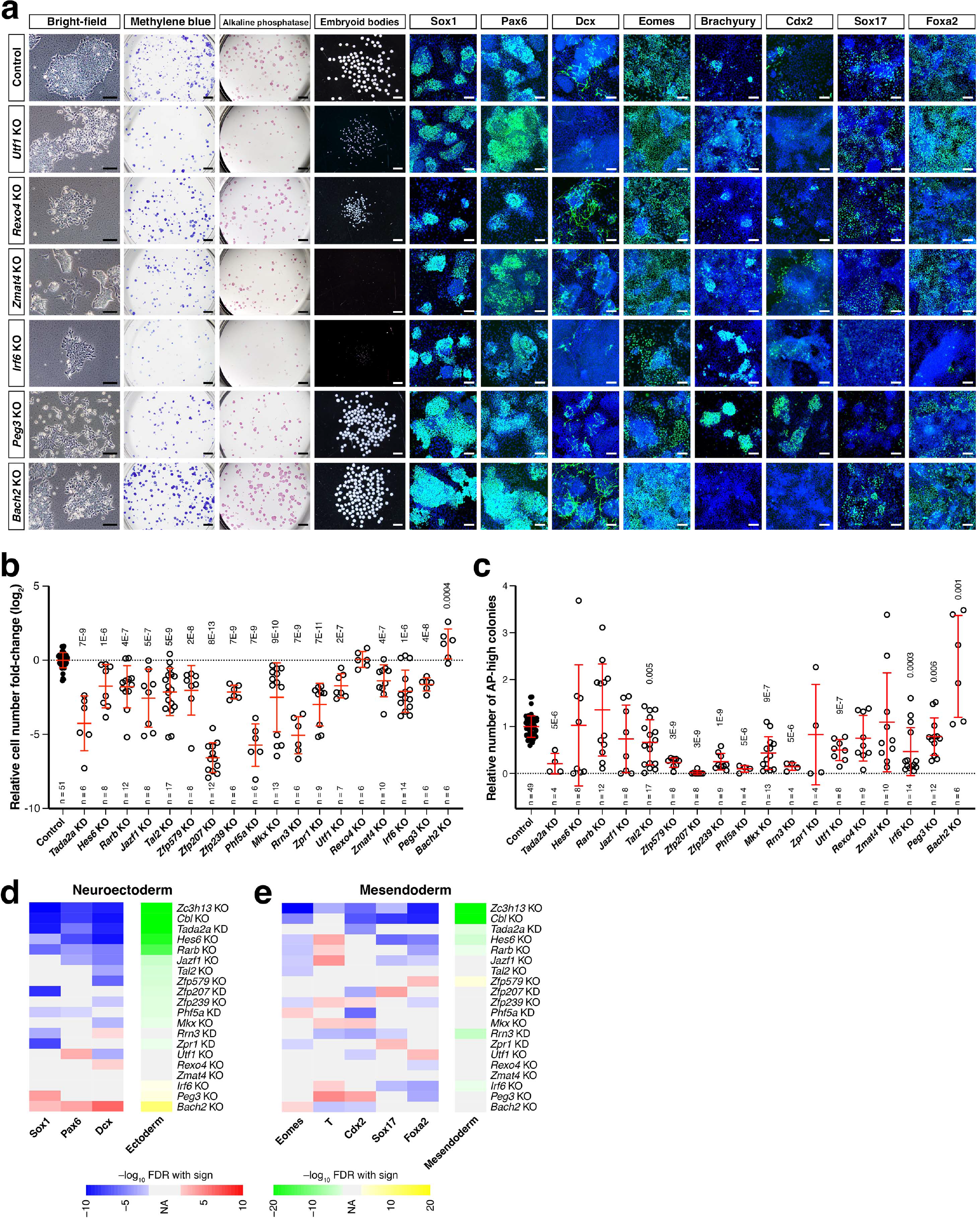
Functional analysis of master regulators (part 2). (a) Analysis of a control EpiSC line and 6 CRISPR KO or shRNA-silenced (KD) lines in self-renewal and differentiation assays. Data for the control are the same as in Extended Data Figure 3. Scale bars: 100 microns. (b) Fold-change in cell number for KO and KD lines relative to control in colony formation assays, calculated at day 5 relative to the starting cell number at day 0. The fold changes in the KO and KD group were then normalized to the control and log_2_ transformed. Line and whisker plots show the mean ± s.d.; statistical significance is shown as FDR (2-tailed U-test). (c) Quantitation of alkaline phosphatase-staining colonies in KO/KD lines relative to control. Line and whisker plots show the mean ± s.d.; statistical significance is shown as FDR (2-tailed U-test). (d,e) Heatmaps of neuroectoderm (d) and mesoderm (e) marker expression. Statistical significance for the differential protein levels (2-tailed U-test) is shown as -log_10_(FDR). Differential expression (z-scores) across all marker proteins were integrated using the Stouffer’s method and shown in the right-most heatmaps as -log_10_(FDR).

**Extended Data Figure 5:**
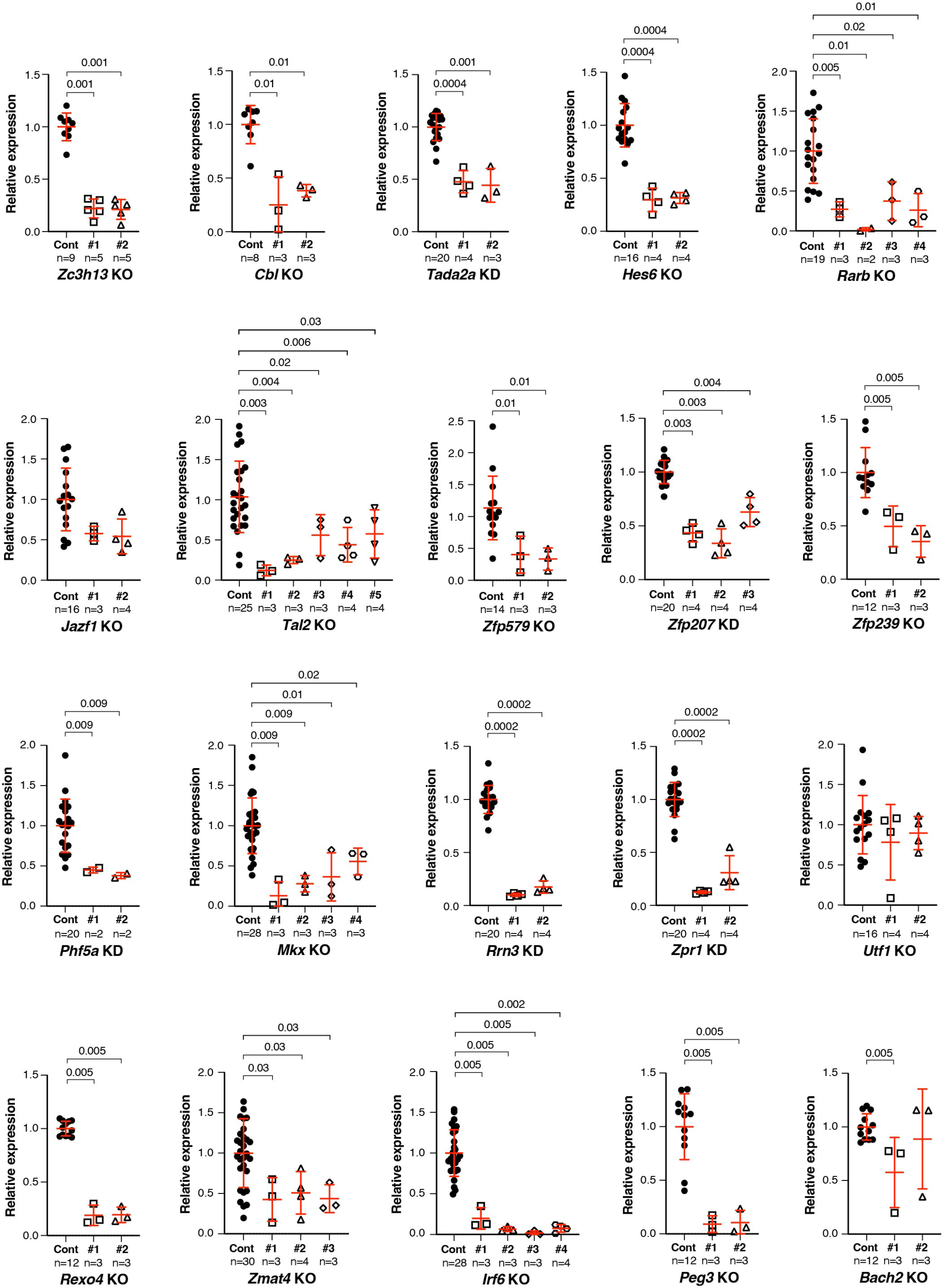
Expression of MRs in CRISPR/Cas9 targeted or shRNA silenced EpiSC lines. Line and whiskers plots show the mean ± standard deviation. Each dot represents a biological replicate. Statistical significance is shown as FDR, 2-tailed U-test for significant (FDR < 0.05) comparisons.

**Extended Data Figure 6:**
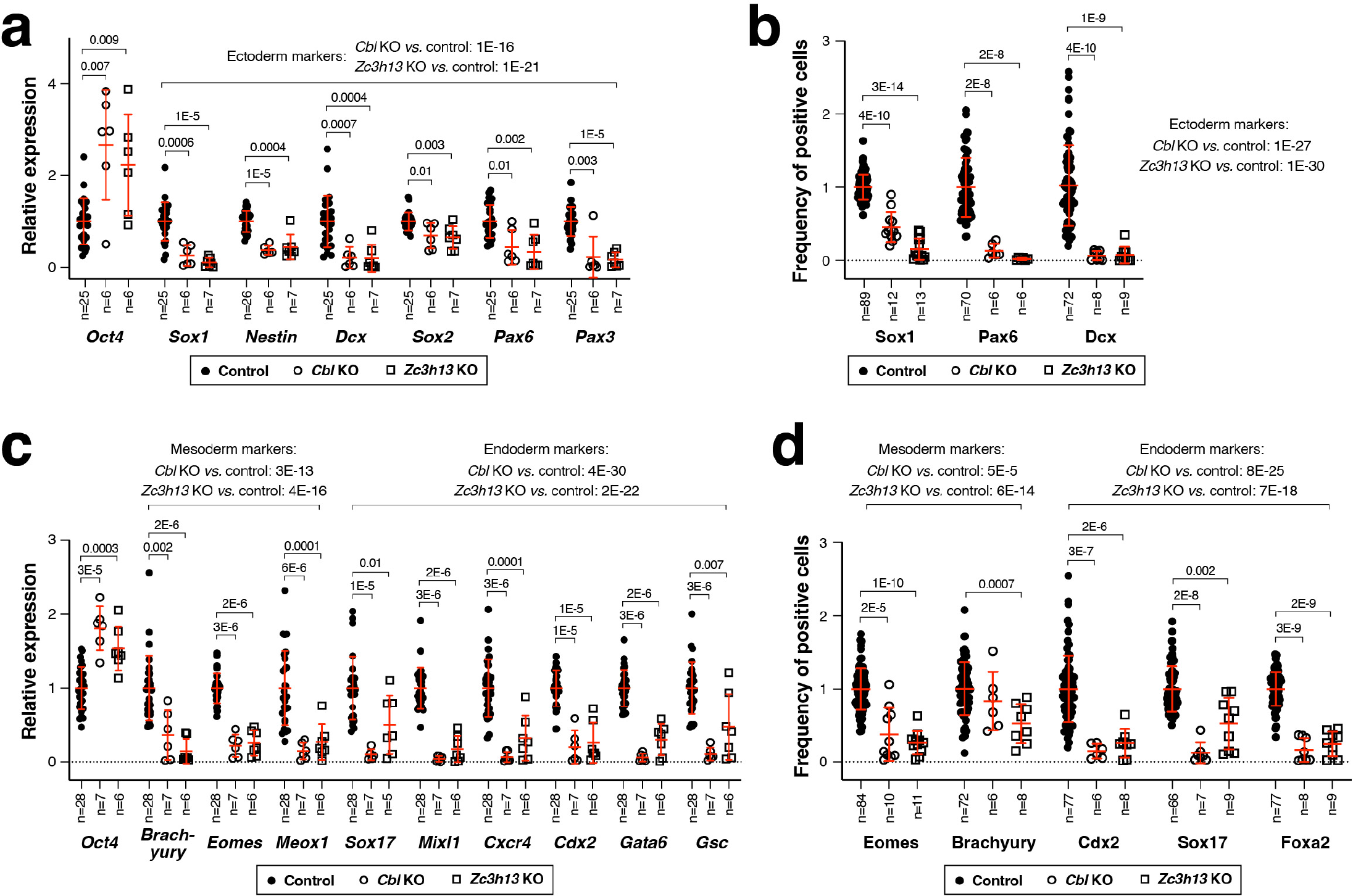
Marker analysis of *Cbl* and *Zc3h13* KO cell line differentiation. (a, c) Expression of marker genes in Cbl and Zc3h13 KO lines relative to batch-matched control EpiSCs after neuroectoderm (a), and mesoderm and endoderm (c) differentiation. (b, d) Percentage of cells expressing marker proteins in Cbl and Zc3h13 KO lines relative to batch-matched control EpiSCs after neuroectoderm (b), and mesoderm and endoderm (d) differentiation treatments. Line and whiskers show the mean ± s.d. for replicates in at least 3 independent experiments. Statistical significance is shown as FDR, 2-tailed U-test for significant (FDR < 0.05) comparisons. The z- scores for markers of the same lineage were Stouffer integrated and the corresponding p-values were corrected for multiple hypothesis testing (B&H FDR).

**Extended Data Figure 7:**
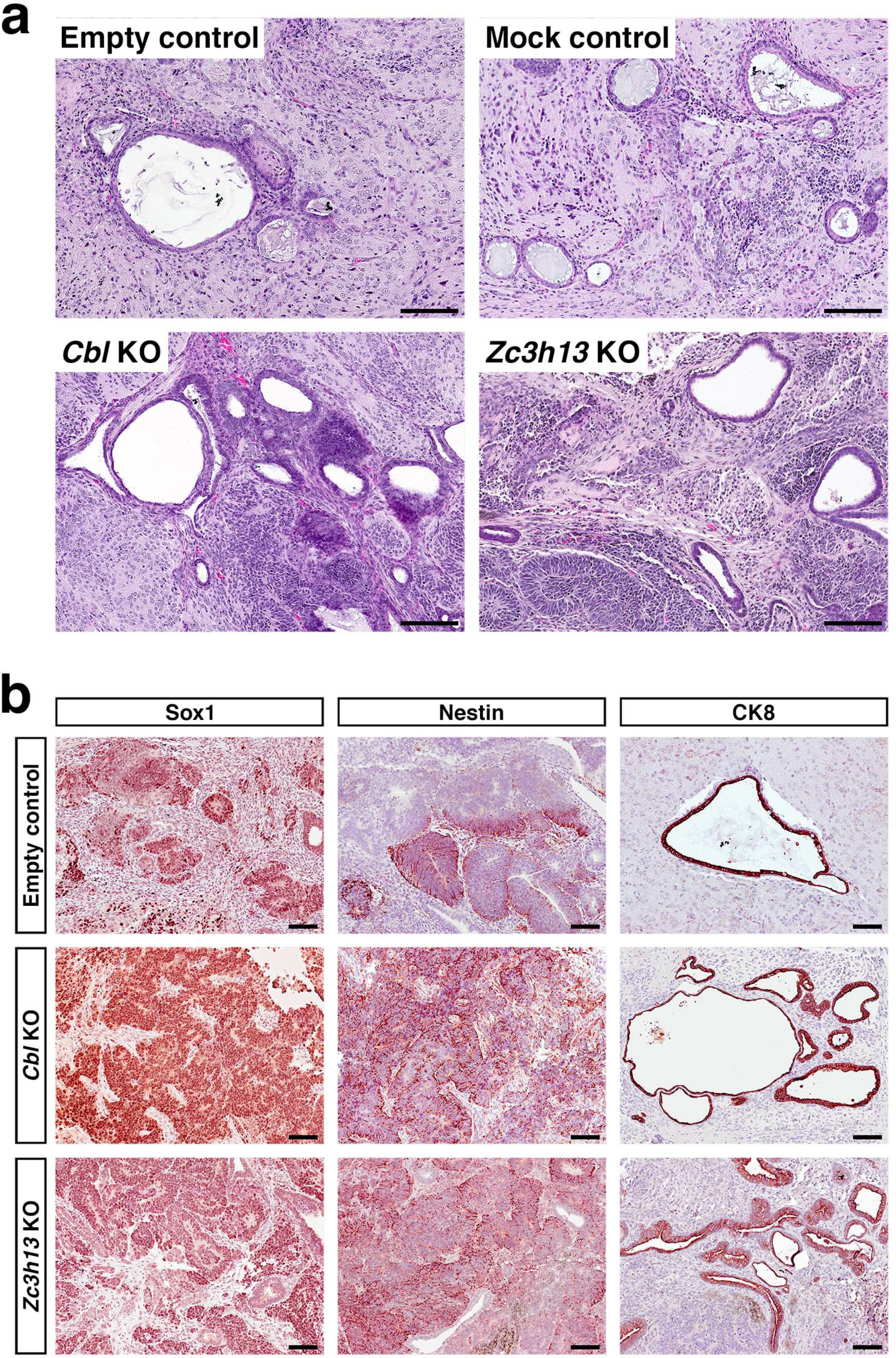
Histopathology of *Cbl* and *Zc3h13* KO teratomas. (a) Hematoxylin and eosin (H&E) staining of representative sections from CRISPR KO and control teratomas. (b) Immunohistochemical staining for neural progenitor (Sox1 and Nestin) and endoderm epithelial (CDK8) markers. Scale bars: 100 microns.

**Extended Data Figure 8:**
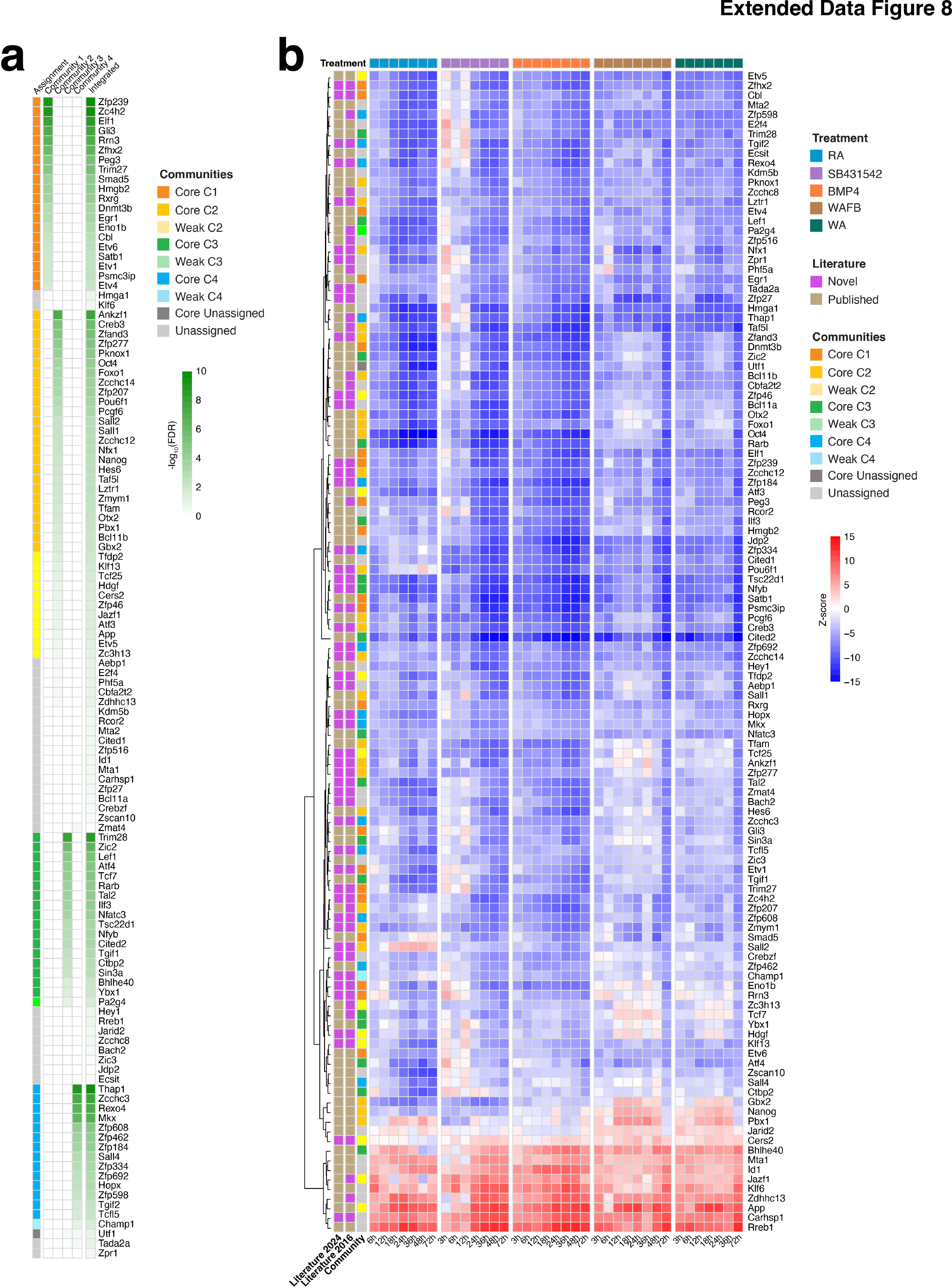
Community membership and relative activity of MR proteins. (a) Heatmap showing the community membership score for the 120 MRs shown in Fig. 3a, calculated by a 1-tailed FET and expressed as -log_10_(FDR). MRs with strong community assignment (FDR < 0.01), MRs with weak community assignment (0.01 ≤ FDR < 0.1), and MRs with non-significant community assignment (FDR ≥ 0.1) are shown by the indicated colors. (b) Heatmap of MR protein activity during the five differentiation time courses quantified using the experimentally curated interactome (Supplementary Table 3.2). Hierarchical clustering was performed using Euclidean distance and complete linkage.

**Extended Data Figure 9:**
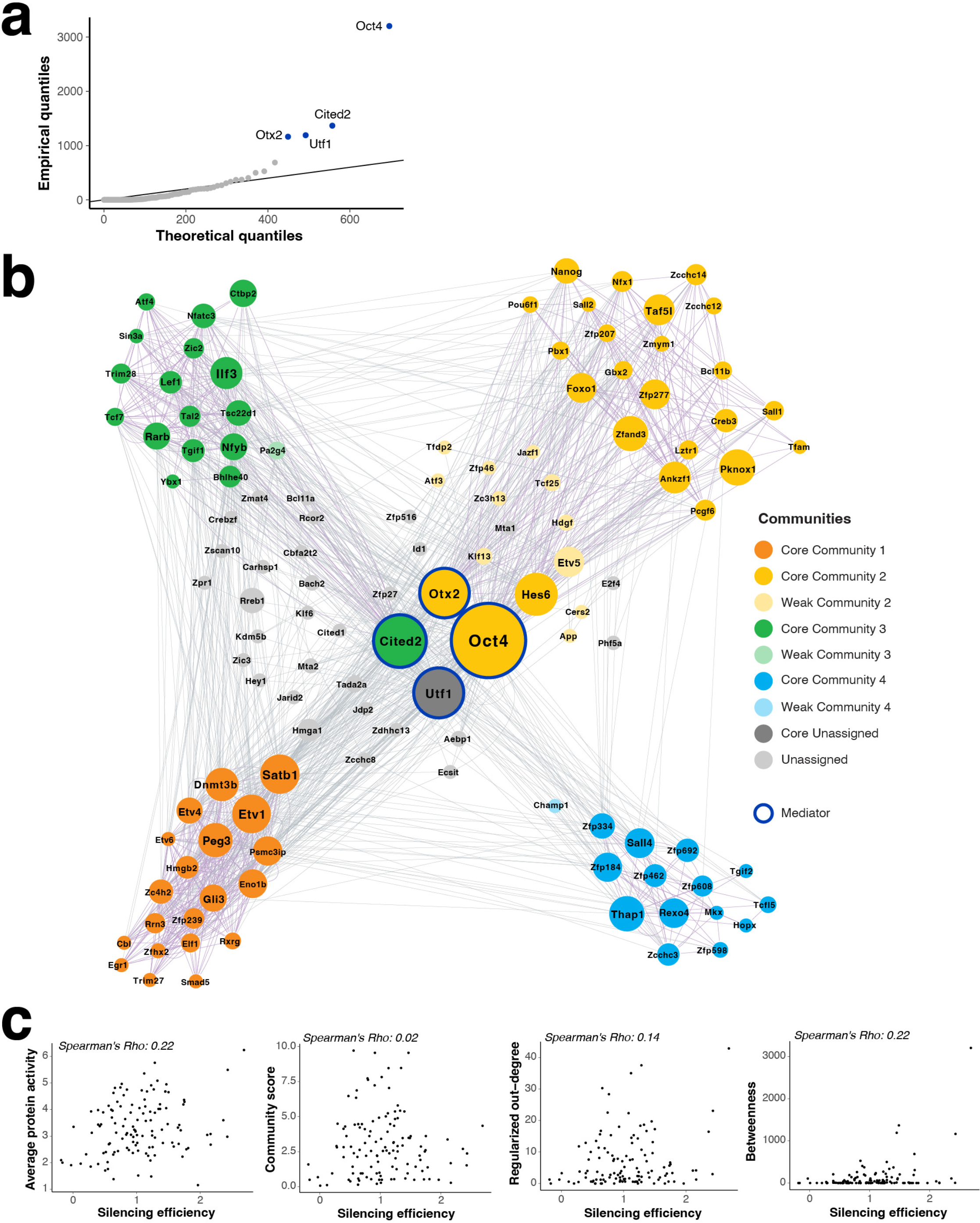
Structure and centrality of the regulatory network for primed state pluripotency. (a) QQ-plot for the theoretical (based on a negative binomial distribution fitted to the data using a maximum likelihood method) vs. empirical quantiles for the Betweenness score. Outlier MRs (nodes with Betweenness score higher than expected given the probabilistic model, shown by the black line, FDR < 0.05) are highlighted in the plot and considered as Mediators. (b) Regulatory network of primed state pluripotency with the size of each node proportional to its betweenness score; colors indicate community assignment (strong, FDR < 0.01; weak, 0.01< FDR < 0.1; unassigned FDR> 0.1). Only significant effects (FDR < 10^-10^, 2-tailed aREA test) are included. Four outlier MRs showing the highest betweenness score (Mediators) are highlighted; Utf1 is shown in dark grey due its high betweenness score and no community assignment. (c-f) Scatter-plots showing the association between shRNA silencing efficiency (x-axis) and average protein activity (c), community score (d), ROD (e), and Betweenness (f). Correlation coefficients (Spearman’s Rho) are shown for each plot.

